# Multiscale Multimodal Characterization and Simulation of Structural Alterations in Failed Bioprosthetic Heart Valves

**DOI:** 10.1101/2023.02.26.529530

**Authors:** Elena Tsolaki, Pascal Corso, Robert Zboray, Jonathan Avaro, Christian Appel, Marianne Liebi, Sergio Bertazzo, Paul Philipp Heinisch, Thierry Carrel, Dominik Obrist, Inge K. Herrmann

## Abstract

Calcific degeneration is the most frequent type of heart valve failure, with rising incidence due to the ageing population. The gold standard treatment to date is valve replacement. Unfortunately, calcification oftentimes re-occurs in bioprosthetic substitutes, with the governing processes remaining poorly understood. Here, we present a multiscale, multimodal analysis of disturbances and extensive mineralisation of the collagen network in failed bioprosthetic bovine pericardium valve explants with full histoanatomical context. In addition to highly abundant mineralized collagen fibres and fibrils, calcified micron-sized particles previously discovered in native valves were also prevalent on the aortic as well as the ventricular surface of bioprosthetic valves. The two mineral types (fibers and particles) were detectable even in early-stage mineralisation, prior to any macroscopic calcification. Based on multiscale multimodal characterisation and high-fidelity simulations, we demonstrate that mineral occurrence coincides with regions exposed to high haemodynamic and biomechanical indicators. These insights obtained by multiscale analysis of failed bioprosthetic valves may serve as groundwork for the evidence-based development of more durable alternatives.

**Graphical Abstract:** 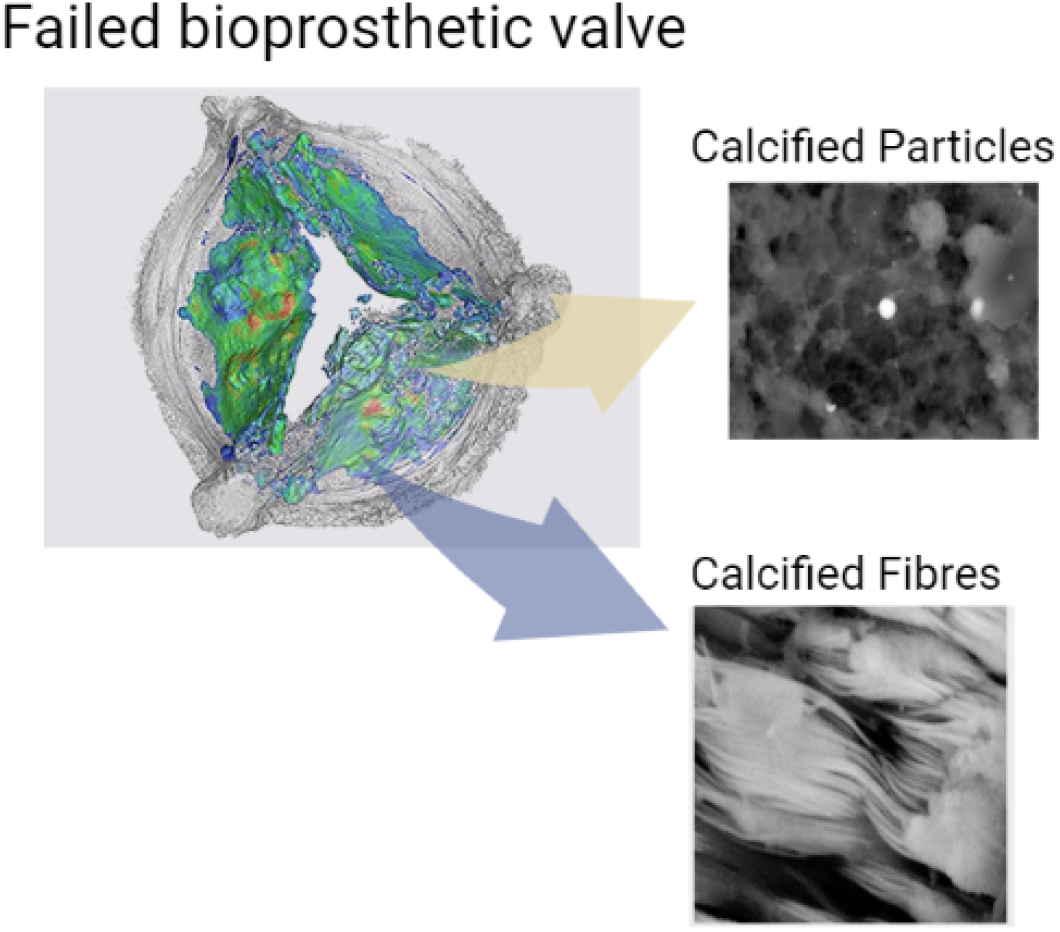

## Introduction

Aortic valve stenosis is the most common valvular heart disease, affecting 12% of the population over the age of 75 (1, 2). The disease involves fibrosis and deposition of calcium phosphate minerals on the valve leaflets, resulting in reduced leaflet movement, blood flow restriction, and other adverse outcomes, including angina and heart failure (3). In moderate to severe aortic stenosis, valve replacement becomes inevitable, usually requiring open-heart surgery in younger and transcatheter techniques in older patients (4). The valve is replaced either by a bioprosthetic (xenografts, allografts or exceptionally homografts) or a mechanical heart valve (4). Unfortunately, subsequent calcification in bioprosthetic valves is frequently observed, limiting prosthesis durability significantly. Although mechanical valves overcome this drawback, they require lifelong anticoagulation therapy and put patients at risk of bleeding and thromboembolic complications (4). Thus, they are primarily used in younger patients who do not qualify for bioprosthetic valves due to their limited durability.

Understanding the role and origin of calcification in native heart valves has been a major topic of research. Medical imaging studies using x-ray computer tomography and echocardiography have provided insights into the critical role of calcification in aortic valve stenosis, as increasing amounts of calcification correlate with disease severity in most patients (5, 6). Studies have suggested that biomechanical stresses contribute to the development and progression of aortic valve stenosis (7–9). For example, the earlier calcification onset observed in the non-coronary leaflet has been thought to be related to higher oscillatory shear stresses (7, 8). In addition, considerable efforts by the molecular biology research community have focused on the mineralisation mechanisms in native valves, suggesting a complex, multifaceted process involving osteoblastic differentiation, downregulation of anti-calcific factors, mechanodynamics, shear stresses, chronic inflammation, as well as lipid accumulation (3, 10, 11).

Despite the growing mechanistic understanding of native heart valve calcification, it remains unclear whether similar mechanisms trigger the development of minerals in bioprosthetic valves. While some studies report shared aetiology (12), different cell types and protein contents associated with the mineral in native versus bioprosthetic valves have been reported (13, 14). Calcification in bioprosthetic valves and subsequent failure have also been widely associated with the pretreatments used prior to implantation, i.e. tissue damages induced by preservation (3, 15, 16). Recent research has indicated that calcification could result from an unmitigated immune response due to insufficient glutaraldehyde fixation (very low concentrations are usually used) (17). Consequently, research on masking antigenicity through stronger fixation and decellularisation has been initiated (18, 19). An overactive immune system is also believed to be the reason young patients undergoing bioprosthetic valve replacement are more prone to early calcification (20, 21). Finally, the contribution of biomechanical forces is debated, as it is suggested that areas experiencing higher biomechanical stress are subject to higher incidents of structural alterations (22, 23).

Even though considerable advances have been made, bioprosthetic valvular failure remains a significant and unresolved issue (14). The apparent complexity of the mineralisation mechanisms has highlighted the need for a more in-depth, systematic understanding of the mineral structures present in bioprosthetic tissues. This is best illustrated by thorough electron microscopic investigations, which lead to significant advances in native valve calcification research, identifying distinct structures, including; calcified particles and compact calcification (2, 24, 25). Similar structures have also been reported in explanted bioprosthetic valves; however, lacking histoanatomical context and high-resolution analysis (2, 26). This disconnect between the clinicopathological investigations and the nanoanalytical findings obtainable by electron microscopy is prohibitive to a holistic understanding of bioprosthetic valve failure (21).

In this work, we combine clinically established methods such as (micro) computed tomography and histological analysis with cutting-edge material sciences methods, including scanning small angle x-ray scattering and electron microscopy, to provide comprehensive insights into mineralisation and structural changes in failed surgical bioprosthetic valve explants across scales. The findings are spatially correlated to macroscopic information on the biomechanical forces acting on the leaflets obtained through numerical simulations of the fluid-structure interaction between valve tissue and blood flow. Our analysis uncovers structural alterations in the collagen organisation associated with fibrillar mineralisation and provides a comprehensive characterisation of the minerals found in failed bioprosthetic valves with full histoanatomical context. Furthermore, it is shown that the spatial distribution of these minerals can be associated with regions exposed to complex blood flow patterns characterised by increased oscillatory shear index, topological shear variation index and residence time.

## Materials and Methods

### Sample collection

Failed bioprosthetic heart valves (Table S1) and native aortic valves (analysed as a reference) were kindly provided by the Department of Cardiovascular Surgery of the University Hospital Bern (Inselspital), Switzerland, following the obtainment of general written consent. All bioprosthetic valves were Perimount Magna Ease, tricuspid valves made of pretreated, fixed, bovine pericardium. Failed explanted valves were shipped in ethanol, post-fixed using 4% of paraformaldehyde (PFA) for 24 hours and then stored at 4 °C until further analysis. A new bioprosthetic valve (bovine pericardium Perimount Magna EASE 23 mm; stored in glutaraldehyde) was also obtained to serve as a control for the presence of minerals prior to implantation, and two old expired bioprosthetic bovine pericardium valves (Perimount Magna EASE 23 mm and 21 mm stored in glutaraldehyde) were used as a control for the induction of calcification during storage.

### Micro X-ray computed tomography (CT)

For global microCT measurements, the valves were removed from PFA and placed in a saturated water atmosphere. The samples were mounted on a sample holder and scanned on a cone-beam RX Solutions Easy Tom XL microCT system, with a flat panel Varian PaxScan detector operated at an accelerating voltage of 140 kV with a tube current of 180 μA. The beam was pre-hardened using two copper filters (width: 0.7 mm) to reduce metal artefacts originating from the bioprosthetic valves’ metal frame. All valves were scanned in the same manner. The voxel size of the microCT scans was around 20 μm. Manipulation of the data to further reduce metal artefacts (metal segmentation, histogram in-painting, and image fusion) was done using the RX solutions software. Following, leaflets of interest were cut out for higher resolution, local scans to be acquired. Valve leaflets were dissected using standard dissection tools and were placed in a saturated water atmosphere. High-resolution scans were obtained using an accelerating voltage of 75 kV and a tube current of 200 μm (for a global CT – voxel size ~ 14 μm) and 150 μm (for a local CT – voxel size ~ 3 μm). Prehardening of the beam was also performed using a 0.5 mm aluminium filter. The 3D segmentation, rendering and analysis were done with VG Studio and FIJI ImageJ. For the cumulative spatial distribution of calcification, the images were registered using landmarks and transformed into binary text files where 1 corresponded to black (calcification) and 0 to white (tissue) and added up to evaluate the areas where most leaflets presented calcification.

### Scanning electron microscopy

Regions of interest from isolated leaflets were identified based on the microCT data and were cut out using standard dissection tools for scanning electron microscopy (SEM). Samples were then dehydrated using an ethanol gradient of increasing concentrations, left to air dry, and mounted on silver stubs with carbon paste (Plano Leit C). The samples were silver painted and carbon-coated with 10 - 15 nm of carbon using a Safematic CCU-010 coater. An FEI Quanta 650 field emission gun environmental scanning electron microscope was used for imaging. Secondary electron (SE) and backscattered electron (BSE) signals were collected. Energy Dispersive X-ray (EDX) Spectroscopy analysis was carried out using the Thermo Fischer Pathfinder SDD EDX system. An accelerating voltage of 10 kV was used for all imaging and analysis. Size distribution analysis was carried out using FIJI ImageJ software. Box plots show interquartile (IQR) range and average values, and the whiskers indicate the minimum and maximum values.

### Histology

A selection of isolated leaflets (based on the degree of calcification) was sent to SophistoLab, Muttenz, Switzerland, for paraffin embedding and sectioning. Sections were stained with hematoxylin and eosin (H&E) to observe the tissue architecture. The presence of calcification (appearing as brown/black) was examined through Von Kossa staining. The collagen network was also visualised through PicroSirius red staining and polarised light microscopy, which enables the visualisation of the heterogeneity and organisation of collagen fibres. All images were recorded on a Zeiss AxioImager.Z2 using the brightfield or darkfield mode and the 5x, 20x objectives.

### Confocal and two-photon microscopy

Collagen organisation and correlation with the mineral were evaluated using a Multi-Photon Leica SP8 MP microscope. Standard histological sections were used. The samples were incubated with a dilution of 1:10, near-infrared bisphosphonate-based calcium dye (OsteoSense 680, Perkin Elmer) for 20 minutes, washed with PBS and mounted using Fluoroshield mounting medium. The OsteoSense stained calcification was visualised using confocal imaging and collagen fibres through second harmonic generation imaging, using a MaiTai XF laser tuned at 880 nm and collecting the emission signal at 440 nm. Two internal and two external (a forward and a backward) detectors were used to capture the maximum collagen orientations. All images were captured using the 25x objective.

### Small angle X-ray scattering

Scanning SAXS was performed at the cSAXS beamline at the Paul Scherrer Insitute with a monochromatic synchrotron-based x-ray beam focused to 25 x 25 μm^2^ and a fixed-exit double Si(111) monochromators at an energy of 11.2 keV. 2D scattering patterns were recorded on a Pilatus 2M detector at two sample-detector distances, 7 m and 2 m, respectively, calibrated from the scattering pattern of silver behenate. The transmitted beam intensity was measured with a photodiode placed on the beamstop inside the steel flight tube under vacuum conditions. Bioprosthetic heart valves were measured in Kapton pockets filled with PBS buffer to maintain their hydration level. All three samples were raster-scanned at 100 x 100 μm^2^ at a detector distance of 2 m to get an overview of the full leaflets. High-resolution scans for selected regions were performed at 50 x 50 μm^2^ and a sample-detector distance of 7 m to complement the overview scans and assess potential changes in the collagen fibril diameter of non-implanted and explanted calcified heart valves. For the calcified sample, the selected region of 3 x 5 mm^2^ contained a combination of calcified and un-calcified tissues. A 1 x 1 mm^2^ region was scanned for the non-implanted leaflet. Radial integration of 2D scattering patterns was performed following the azimuthal segment procedure using the cSAXS MatLab analysis package (Scanning SAXS Software Package, https://www.psi.ch/en/sls/csaxs/software (accessed: April 2018)). Radiation damage check was performed similarly for all samples prior 2D scans by monitoring a change of scattering when exposed to repeated 0.1 s exposure time over several seconds. The collagen fibril diameter was estimated by the peak maximal of background subtracted Gaussian fit of the fibrils in bundles’ d-spacing peak between 0.02 < q < 0.09 nm-1 (Fig. 5). Orientation analysis of collagen fibril diameter peak and tropocollagen 5^th^ order of the diffraction peak stemming from the staggered supramolecular structure along the collagen fibre was also done. The 5th order peak was selected to avoid overlap with scattering from d-spacing of fibres in bundle (aka “collagen diameter”) higher harmonics. A q-range selected between the 4^th^ and 5^th^ collagen peaks at 0.417 < q < 0.450 nm^-1^ was selected to represent the orientation of the collagen or, when present, the mineral phases nanostructures (referred to as collagen/mineral *q*-equivalent domains (CMQED) within a size range of 13 < d < 37 nm. Orientation analysis was performed on these regions following the analysis routine established by Bunk et al. 2009 (27). The colour wheel contains information on the main orientation of nanoscale features (hue), their degree of orientation (saturation) and the amount of scattering material. With this encoding function, a material with a low electron density contrast will appear black, dense material with low preferential orientation will appear with a light colouration depending on the angle, and dense material with a strong preferential orientation will appear with vivid colours. Comparison of the three different scattering domains considered in this study, the d-spacing peak of collagen fibres in bundles, the 5^th^ collagen peak and the CMQED allow us to probe, the average collagen bundles’ main orientation, the average orientation of the tropocollagen molecules within a fibril, and the orientation of the mineral phases forming within the collagen structure respectively.

### Coupled simulation of blood flow and leaflet dynamics

The computational method for the high-fidelity simulation of the blood flow and the mechanical dynamics of the leaflets relies on a fluid-structure interaction (FSI) approach based on an immersed boundary method (28). The incompressible Navier-Stokes equation is solved on a fixed staggered Cartesian grid of about 4 million cells using a six-order finite difference scheme and an explicit third-order Runge-Kutta timestepping scheme for the advective term, and a semi-implicit Crank-Nicolson scheme for temporal discretisation of the diffusive term (29). The elastodynamics equation is solved on a moving tetrahedral mesh (consisting of about 200,000 tetrahedra constituted of 4 nodes each) using the finite-element method with affine finite elements and a Newmark time-stepping scheme(30). The strong coupling of these two sets of equations is based on a parallel variational transfer and an iterative procedure to ensure traction continuity and the equality of the solid and fluid velocity fields at the fluid-structure interface (31). A fibre-reinforced model is used to characterise the anisotropic material properties of the leaflets (32, 33), while the material properties of the aortic wall and the supporting crown of the leaflets are described by a linear elastic material model (34). The strain energy density function Ψ_HGO_ used to define the constitutive relation of the (noncalcified) leaflets modelled with a nearly incompressible anisotropic hyperelastic material is given by (32):

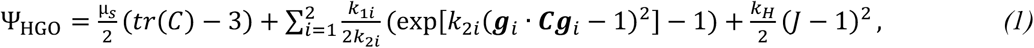

with *μ_s_, k_li_, k_2i_*, the fitted parameters of the model (33), *k_H_*, the penalty coefficient for the incompressibility constraint, **g_i_**, the fibre orientation (two families depending on whether the radial or circumferential direction is considered) and **C**, the right Cauchy-Green strain tensor (see definition in Eq. 7). The geometrical setup is similar to those for models of bioprosthetic heart valves made from bovine pericardium and mounted on a rigid polymeric ring (35). It is worth noting that the considered geometry of sinus of Valsalva in the computational model does not include coronary branches; hence no differentiation between right, left and non-coronary cusp for the nomenclature of the leaflet is provided. Two geometries of valve leaflet are considered to compare the microscopic characterisation of the zones onto the leaflet surface where calcification preferentially occurs. These leaflet geometries have been chosen due to their unstable flutter motion noted in the FSI simulations at peak systole (34, 35). The flow conditions considered in the computational study are peak systolic ones under a pressure drop across the valves and in the ascending aorta of 8 mmHg over a time span of 0.3 s.

### Indicators for the calcification characterisation from FSI simulations

The analysis of the flow field at the fluid-leaflet interface is based on blood-flow-related indicators found in the literature to trigger endothelial cell response at the vascular wall through biochemical signals leading to atherosclerotic lesions (36–39). In this study, we hypothesise that, although these descriptors are commonly used to predict atheroprone regions resulting in changes in the vascular wall morphology, these indicators could also be valuable to macroscopically highlight zones prone to mineral nucleation and growth in bioprosthetic heart valves. A list of five indicators depending on the blood velocity close to the leaflet and their definition are provided below.

The wall shear stress vector of a Newtonian fluid, which is defined as the projection of the viscous stress tensor onto the leaflet wall normal vector is given by (38):

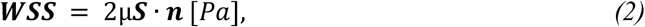

with **S**, the strain-rate tensor of the local fluid element, **n**, the normalised vector normal to the leaflet wall and μ = 0.004 Pa, the dynamic viscosity of blood.

The wall shear stress vector is the quantity based on which the other indicators are computed in the rest of this paper, and the time-averaged value of its magnitude (TAWSS) corresponds to one indicator used in the present study. The time-averaged pressure (TAP) represents the diagonal part of the total stress (the fluid exerts onto the leaflet surface) tensor.

The oscillatory shear index (OSI)) is used to quantify the temporal variability of the wall shear stress vector and is defined as (37):

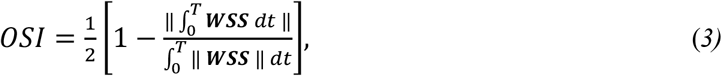

With ||***WSS***||, the L2-norm of the wall shear stress vector and T, the simulated time over peak systole.

The relative residence time (RRT) (36, 37) is calculated from the TAWSS and OSI indicators as follows:

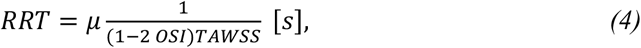

The topological shear variation index (TSVI) as proposed in (36, 40) is considered in this study as an indicator to represent the expansion and contraction at the leaflet surface due to the spatial variation of the wall shear stress vector and calculated by taking the root mean square of the instantaneous temporal fluctuations of the wall shear stress divergence over the systolic phase. This is thus calculated with the following equation:

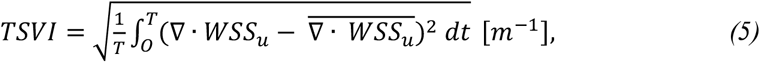

with **∇** ·, the divergence operator, *ō*, time-averaging operation and **WSS_u_**, the wall shear stress unit vector 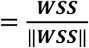.

In order to characterise from the computational fluid-structure analyses the mechanics of the leaflets and the influence thereof on the fibre damage in the tissue-based bioprosthesis, two additional indicators pertaining to the leaflet dynamics are computed. The first one is the time-averaged equivalent tensile strength or von Mises stress, which is dependent on the symmetric Cauchy stress tensor σ:

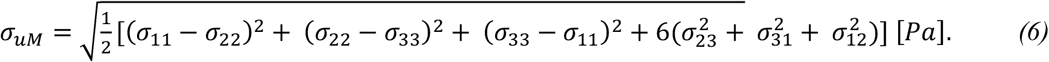

The second indicator is the time-averaged scalar strain defined as the trace of the right Cauchy-Green strain tensor **C**:

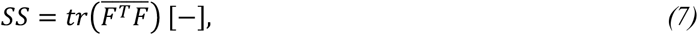

With **F** = ∇_*x*_*u_s_* + **I**, the deformation gradient tensor; **u**_s_, the displacement vector of the leaflet from the initial configuration and **I**, the identity tensor.

### Equation for the reconstructed calcification intensity field

In order to reconstruct a calcification-prone intensity field that depends on the aforementioned indicators, a procedure has been established. Similarly to the approach presented in Corso et al. (41) for establishing fitted models for the evaluation of pressure gradient and haemodynamic stresses, this procedure relies on a non-linear least-square minimisation problem. The non-linear function for which the coefficients need to be fitted can be expressed as follows:

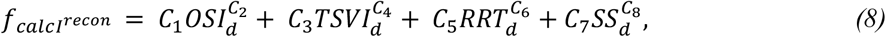

with *C_i_*, the coefficients to be fitted. These coefficients are set to be positive. The choice of the four indicators (namely WSS-based OSI, TSVI and RRT and motion-based SS) for the minimisation procedure stems from quantitative data observed in the results for each individual indicator (see Figure 6), in line with the spatial distribution of the microCT reference data. It is worth noting that the indicators’ values are first normalised so that the range of values for each indicator varies from 0 to 1. In addition, the values of the normalised indicators are distributed over bins of 0.01 and then normalised so that the integral over the discrete distributions is equal to one. The normalised distribution of each indicator is referenced with the subscript “d” in Eq. 8. The calcification intensity observed from the microCT measurements calc**I**^ref^ (cf. Fig. 1) is also first normalised so that the intensity value ranges from 0 to 1 in order to be used as reference data for the minimisation using the least-square method with the objective function 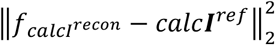.

**Fig. 1.**
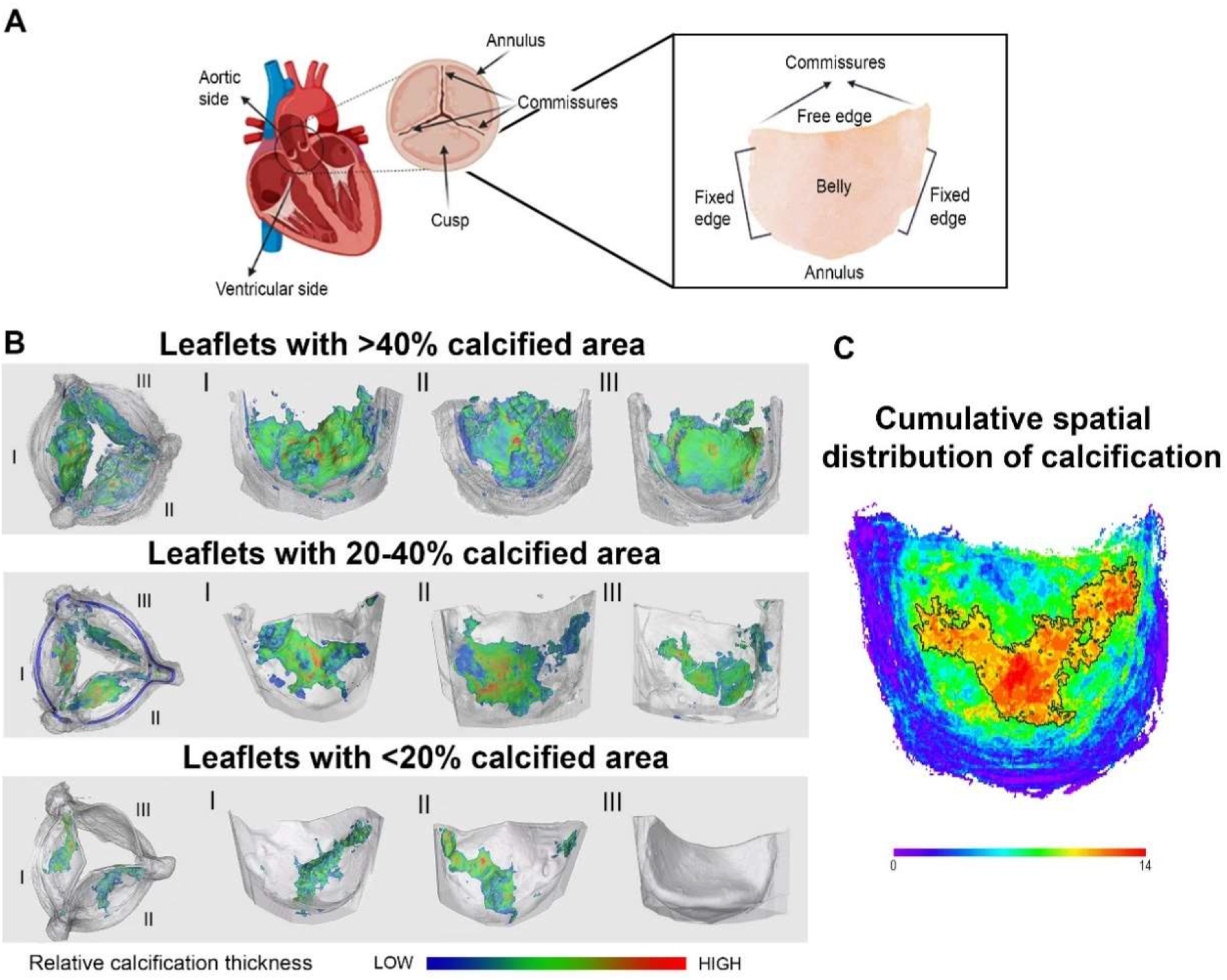
Macroscale anatomical analysis of calcification in bioprosthetic valves. **(A)** Schematic diagram of a tricuspid aortic valve and individual isolated cusp for reference. **(B)** Representative global microCT of calcified bioprosthetic valves, and **I**, **II** and **III** corresponding higher magnification global microCT of individual valvular leaflets presenting different calcification degrees. **(C)** Heatmap of the relative cumulative spatial distribution of calcification using all calcified leaflets (n=14) where purple (intensity 0) and blue correspond to non-calcified areas and red (intensity 14) corresponds to calcified areas observed in all leaflets, indicating that the central parts of the leaflet belly (black outline) was calcified in most (orange areas) or all (red areas) leaflets. Shapes aligned to outer leaflet dimensions.

### Statistical analysis

Origin Pro 2020b software (OriginLabs, Massachusetts, USA) was used for all statistical analyses. Linear regression fitting was carried out to evaluate the relationship between implantation duration and calcification area percentage and maximum calcification thickness (N=21 (all explanted leaflets obtained)) and between maximum thickness and the calcification area percentage (n=14 (all macroscopically calcified leaflets as identified by CT data)). The results are reported using the Pearson’s correlation coefficient (r). The mineral dimension data was plotted in boxplots as the interquartile range (IQR) with whiskers representing the minimum and maximum values, and individual data points included as rhombic points.

## Results

### Macroscale structural assessment shows calcification of the leaflet belly

To assess the presence, amount, and anatomical location of calcification in failed explanted surgical bovine pericardium bioprosthetic valve leaflets (Fig. 1A) at the macroscale, microCT scanning and reconstruction were performed. The twenty-one valve leaflets (isolated from seven explanted heart valves) analysed by CT, contained a variable degree of macroscopic calcification (non-calcified (n=7), calcified (n=14), see Table S1). It was observed that in highly calcified leaflets, the mineral covered the entire belly area affecting the free edge and the commissures (Fig. 1B). In moderately calcified leaflets, the mineral was present in irregular patterns spreading from the lower and central parts of the belly near the annulus to the commissures, with the free edge remaining largely unaffected (Fig. 1B). A similar pattern was observed in the least calcified leaflets with commissures and free edge largely calcification-free (Fig. 1B).

Interestingly, the calcification patterns in the different leaflets indicate that the region most commonly calcified is the central area of the leaflet belly (Fig. 1C, region indicated by black outline) with the area appearing as saturated red (corresponding to intensity value 14) being calcified in all leaflets containing microCT-detectable minerals. This data could indicate a progressive process where calcification occurs initially in the lower and central parts of the belly and extends gradually to the entire leaflet. The presence of a progressive process is further supported by strong correlations (Pearson’s r > 0.89) between the amount of leaflet calcification and maximum mineral thickness (growing up to 1.33 mm) with the duration of implantation (Fig. S1). However, despite the observed trend of mineral accumulation in the central part of the belly, the variation in mineral patterns in the peripheral regions varies significantly, possibly due to a complex, multifaceted formation mechanism.

### Micro and nanoscale characterisation reveals the presence of mineralised collagen fibrils

For more detailed information on the mineral structures, high-resolution microCT reconstructions (voxel size of 3 μm) of individual leaflets were carried out, revealing the presence of fibrous structures (Fig. 2A and S2) along with regions of compact calcification (Fig. 2A III top and bottom). Calcified regions were microdissected from the belly of nine macroscopically calcified leaflets isolated from seven valves and characterised by high-resolution SEM (Fig. 2B). The highly abundant calcified fibres exhibited diameters between 0.5 – 6.4 μm (average: 1.8 μm, IQR: 0.9 μm), were partially formed by organic material based on electron density (Fig. 2B, C and S3) and were observed in all macroscopically calcified leaflets analysed by SEM. Interestingly, these large calcified fibres were formed by bundles of smaller calcified fibrils (average: 109 nm, IQR: 53 nm) (Fig. 2B, C and S3) aligned to some extent along the same axis. Their characteristic diameters match the diameters of collagen type I fibrils (10 - 500 nm in diameter, also observed through SAXS measurements, *vide infra)* and fibres (1 - 20 μm in diameter) (42, 43), thus supporting the widely accepted collagen mineralization theory described for native valves (44). These mineralised fibre and fibril structures were not detected in the native heart valves investigated by us (Fig. S4) nor in previous electron microscopy studies (2, 24, 25). More recently, Gourgas et al. also reported fibrous structures that appear (partially) mineralized in native human heart valves, however, with seemingly low abundance and of unknown origin.

**Fig. 2.**
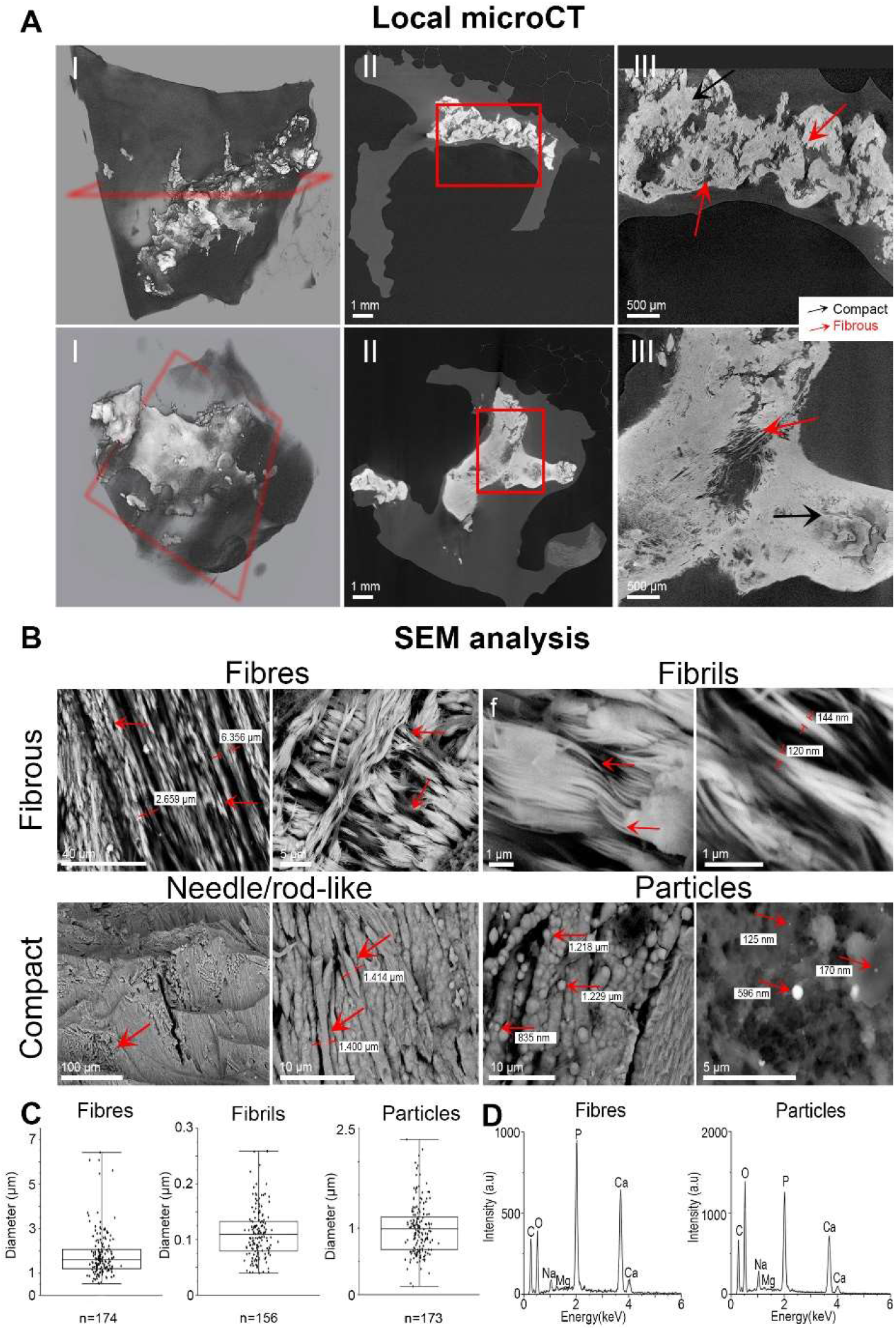
High-resolution imaging of minerals within macroscopically calcified bioprosthetic cusp tissue. Inorganic material is shown as white/light grey and organic material as dark grey/black. **(A) I**, Representative high-resolution microCT reconstructions of valve leaflets with different calcification percentages, 16% and 33%o (top and bottom row respectively) and **II**, cross-sections (indicated as a red rectangle in **I)** of the leaflets showing the calcification shape and **III**, higher magnification microCT images showing fibrous and compact regions. **(B)** Representative high-resolution BSE micrographs of fibrous regions show calcified fibres and fibrils (arrows) and compact regions formed by highly calcified fibres, needle/rod-like structures and calcified particles (arrows) observed in macroscopically calcified leaflets. Some individual calcified particles (arrows) were observed in less calcified areas. (**C)** Fibre (n=174), fibril (n=156) and particle (n=173) diameter size distributions. (**D)** Representative EDX spectra of calcified fibres and particles obtained from macroscopically calcified leaflets.

Extensively calcified merged fibres were also occasionally detected within the more compact mineral structures observed (Fig. 2B), resembling calcified rods and needle-like crystals (Fig. 2B and S3). Additionally, calcified particles (average diameter: 1.0 μm, IQR: 0.5 μm) were detected in all macroscopically calcified leaflet samples (Fig. 2B-D and 4D). These particles often appeared aggregated in highly calcified regions, sometimes forming larger pearl-string structures (Fig. 2B and S3) and were also detected as individual entities in tissue parts where large calcifications were absent (Fig. 2B and S3). Similar particles have also been identified in native valves (Fig. S4), well in line with earlier works (2, 24, 25). Based on EDX analysis, all minerals observed in highly calcified regions were formed by calcium phosphate and contained magnesium in some cases (Fig. 2D and S5).

To distinguish between superficial and deep-lying materials and their characteristics, regions from the aortic and ventricular surfaces of the nine macroscopically calcified leaflets and four non macroscopically calcified leaflets were further analysed by electron microscopy. Calcified particles, fibres and compact minerals (Fig. 3A and B, S6) were found on both the aortic and ventricular surfaces. The calcified particles were predominantly detected on the surfaces (irrespectively of anatomical location (aortic or ventricular side)) of all leaflets (Fig. 3C), suggesting a possible mechanism leading to surface deposition. In contrast, the remaining minerals appeared to surface due to structural growth originating from inside the leaflet, as confirmed by microCT data, which showed that calcifications were primarily found in the middle of the leaflet walls rather than the surface. To ensure that none of these structures result from precipitation due to pretreatment or storage procedures, four expired non-implanted leaflets were processed in the same way and presented no minerals (Fig. 4B).

**Fig. 3.**
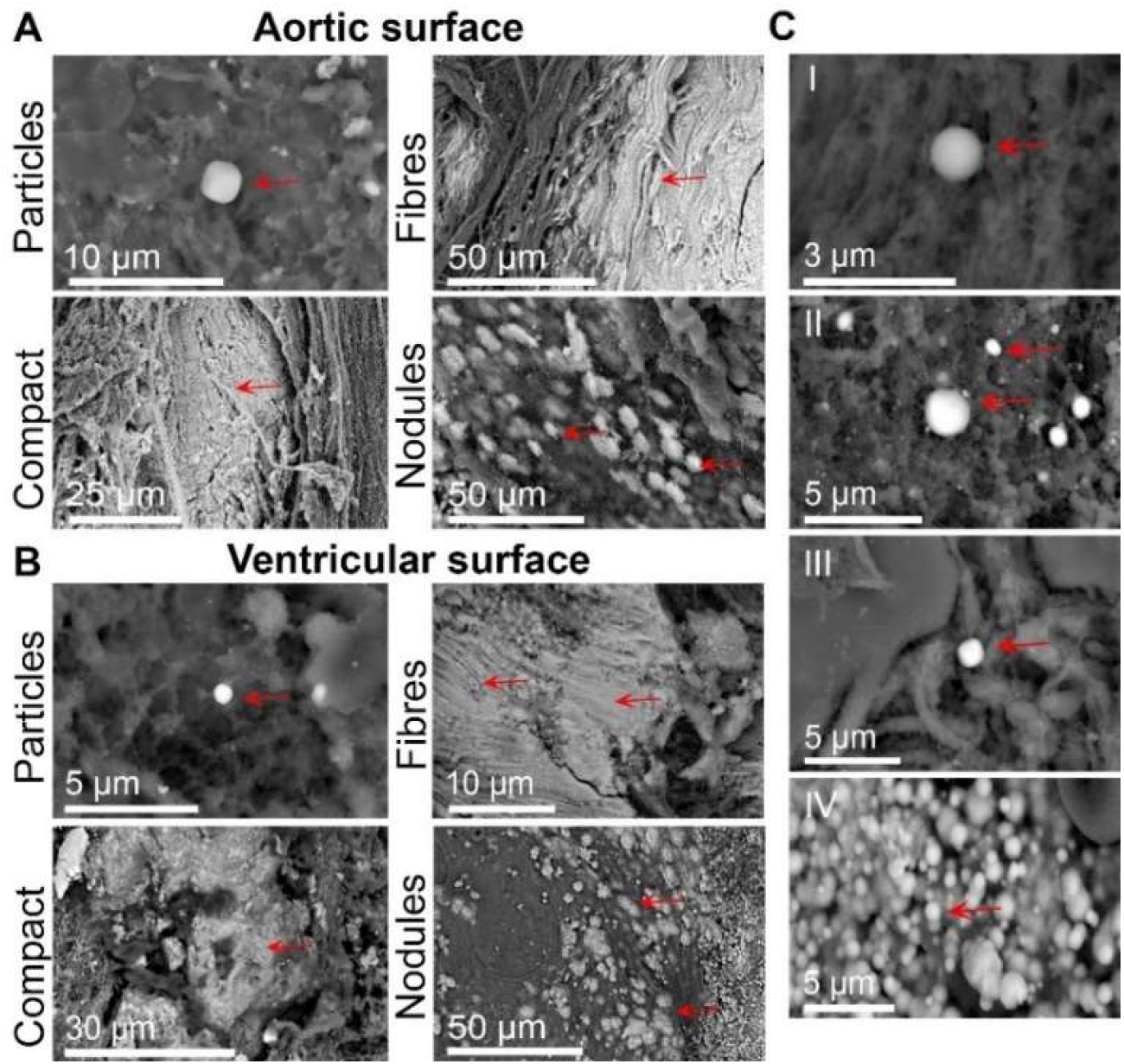
Mineral presence on the aortic and ventricular side surfaces of bioprosthetic cusps. Morphologically different minerals were found; calcified particles, calcified fibres, compact calcification and nodules on the **(A)** aortic and **(B)** ventricular surfaces. **(C)** Calcified particles were observed regardless of the degree of calcification or anatomical location. Scanning electron micrographs of particles found on **I** non-macroscopically calcified, and **II**, **III** and, **IV** macroscopically calcified explanted leaflets of increasing calcification percentages.

**Fig. 4.**
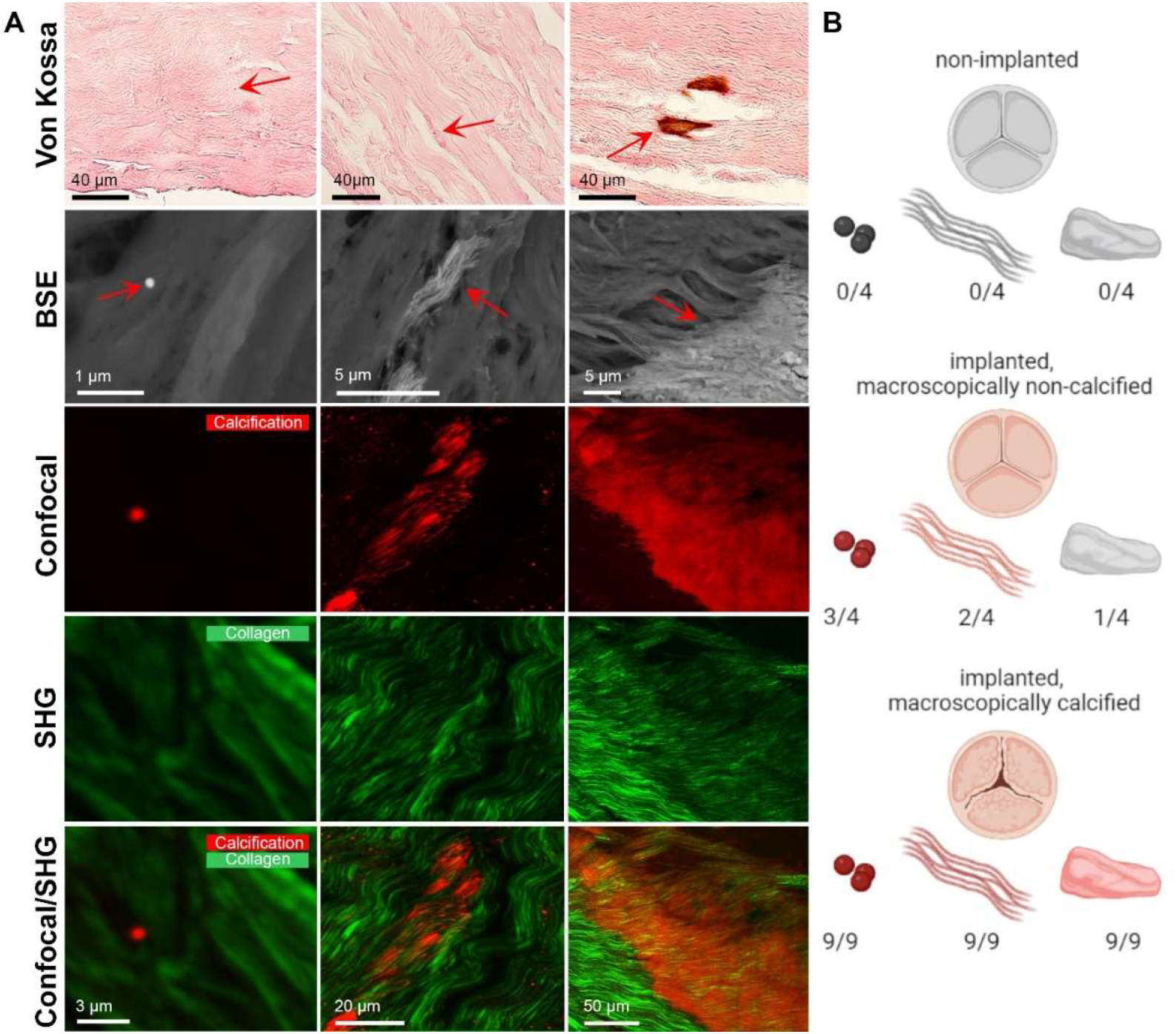
High-resolution imaging analysis of minerals within non-calcified bioprosthetic cusp tissue. **(A)** Calcified regions were observed in Von Kossa stained samples as brown/black spots indicated by arrows. High resolution imaging by scanning electron microscopy enabling morphological and chemical charactierzation of the mineral (BSE, arrows and EDX, see SI Figure S7). Confocal microscopy of the mineral (red), overlaid with SHG data showing the collagen network (green) enables co-localization analysis of the mineral with collagen. Calcified particles, fibres and compact calcifications were identified. **(B**) Prevalence of mineral structures (particles, fibres, compact (macro and micro) calcification) found in non-implanted (n=4), implanted non-macroscopically calcified (n=4) and implanted macroscopically calcified (n=9) leaflets, analysed by micro and nanoscale characterisation techniques.

Thus, it was concluded that the main calcification processes in implanted bioprosthetic valves lead to deposition of calcified particles and their surface and extensive mineralisation of the collagen network. As collagen mineralisation was identified inside the leaflet wall and has not been identified in native valvular leaflets (at very least to no comparable extent), it hints toward a mineralisation process specific to bioprosthetic valvular tissue that initiates within the chemically fixed pericardial tissue. In contrast, the vast presence of the calcified particles both within and on the leaflet wall surface suggests a process of cellular origin (e.g. from macrophages or other cells, cell-derived structures or particles originating from the blood), as previously suggested (45–48) for the same particles (same shape, size and composition) in native valves. Furthermore, similar particles have also been observed in tumour vasculature (49), the Bruch’s membrane of the eye (50, 51), and other cardiovascular tissues(52) suggesting a host-environment origin shared between several pathologies.

### Early-stage mineralisation contains mineralised particles and fibres

To assess the presence of early stages of mineral formation, we investigated non-calcified failed explanted bioprosthetic valve leaflets with high resolution optical and electron microscopy (Fig. 4). Calcified particles were identified (Fig. 4, S7 and S8) through Von Kossa staining and confirmed by nanoanalytical techniques in three out of four non-macroscopically calcified leaflets. Anatomically, they were scarcely observed at various positions on the valve surfaces. EDX analysis indicated that these were always composed of calcium, phosphate and magnesium (Fig. S7).

Backscattered electron microscopy also revealed the scarce presence of calcified fibres and fibrils made of calcium and phosphate (Fig. 4, S7) in half of the investigated non-macroscopically calcified leaflets. Correlative confocal and second harmonic generation (SHG) imaging (Fig. 4A and S8) indicated that these calcified fibres colocalised with collagen fibres, indicating a process of collagen mineralisation, well in line with the SEM investigations (Fig. 2). In contrast, SHG imaging suggests that the calcified particles were entrapped within collagen network gaps rather than deposited onto fibres and were not locally associated with collagen mineralisation (at least at the initial stages). This spatially independent occurrence of the two minerals (calcified collagen fibrils and calcified particles) further supports the existence of two distinct mineralisation phenomena, which may contribute to the creation of larger mineral accumulations (Fig. 2 and 3) over time.

### Microscale characterisation of tissue alterations reveals changes to the collagen structure

Due to the seemingly abundant bioprosthetic tissue-specific collagen mineralisation process, we investigated potential non-calcific tissue alterations that could contribute to its initiation or/and progression. Data acquired from H&E stained sections obtained from the analysed leaflets indicated a loss of cellular components and the thickening of the fibrous network, resulting in more compacted tissue in implanted valves (Fig. S9) compared to non-implanted ones. PicroSirius Red staining further evidence that the collagen network was vastly altered in implanted valves (Fig. S9), even in the absence of calcific deterioration. To gain global information on the collagen network organisation, selected leaflets were investigated by scanning small angle x-ray scattering (SAXS). A new non-implanted bioprosthetic valve leaflet, an explanted macroscopically non-calcified and an explanted macroscopically calcified leaflet were analysed (Fig. 5) (27). A leaflet presenting low degrees of calcification was selected to assess the differences in collagen organisation between calcified and non-calcified regions on the same leaflet. Verifying the histological findings, SAXS analysis indicated differences in the organisation and orientation before and after implantation, even in the absence of macroscopic calcification.

**Fig. 5.**
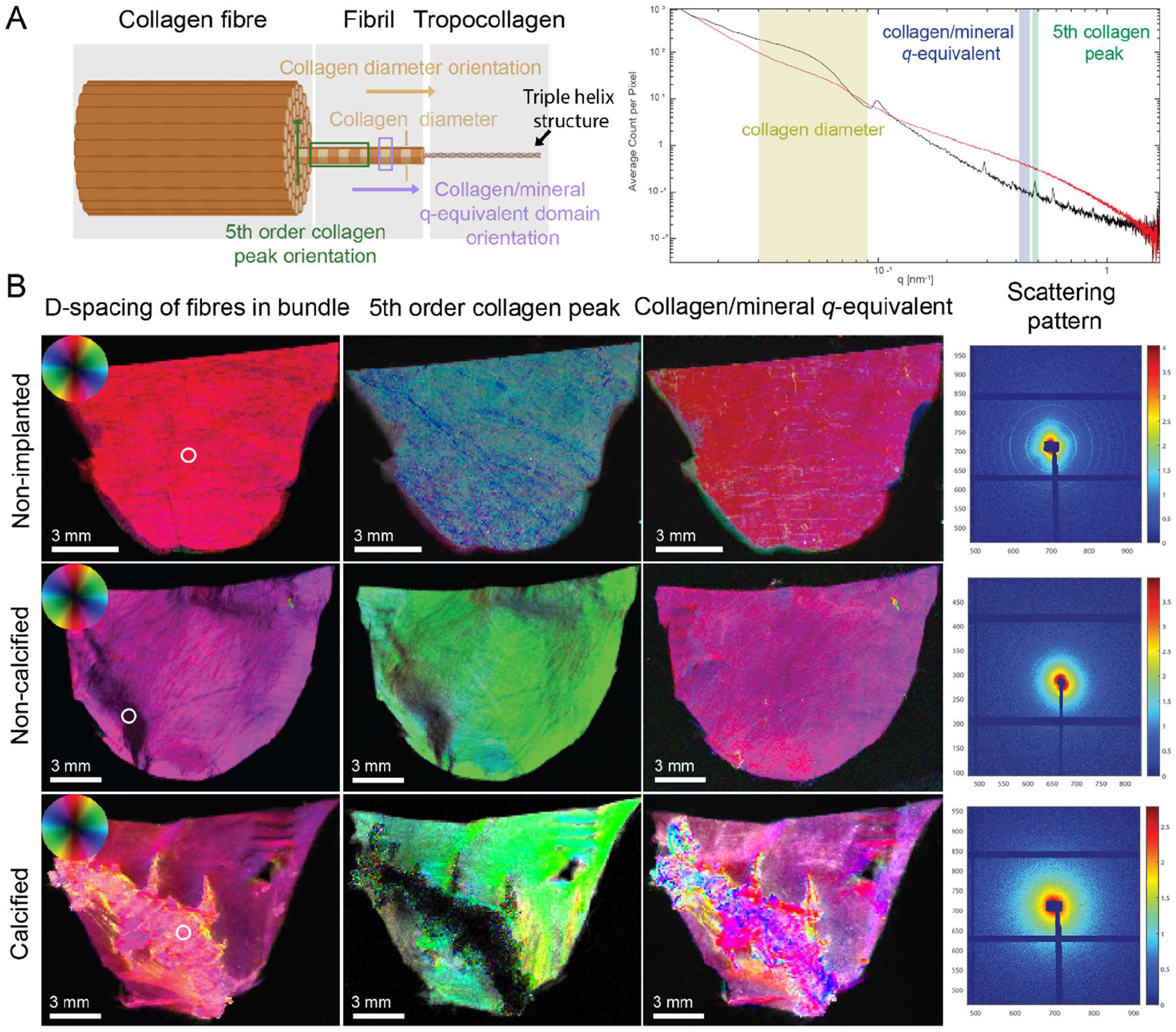
Collagen network SAXS analysis in bioprosthetic valvular leaflets. **(A)** Schematic of collagen structure and diffraction peak plot indicating the collagen diameter, 5^th^ collagen peak and collagen/mineral q-equivalent domain. Black and red curves representing the non-mineralised and mineralised samples respectively **(B)** Orientation analysis of **(I)** the d-spacing peak of fibres in bundles (aka collagen diameter) **(II)** 5^th^ order collagen peak, **(III)** 0.417 < q < 0.450 nm^-1^ collagen/mineral q-equivalent domain and **(IV)** representative scattering patterns of regions marked by a circle, in a non-implanted, a non-calcified explanted and a calcified explanted leaflet. Within this HSV (Hue, Saturation and Value) representation, hue gives information on the orientation according to the colour wheel, saturation on the degree of orientation and the value on the density of the scattering material.

On the one hand, the non-implanted bioprosthetic valve leaflet presented homogeneously a typical fibrillary diameter of 141.46 ± 1.44 nm (Fig. S10) and a scattering pattern characteristic of healthy collagen (Fig. 5 and S11) (‘eye brow’ pattern), known to originate from the triple helix structure of the tropocollagen molecule (53). Similarly, the q-regime corresponding to the d-spacing of the fibril in bundle – corresponding to the collagen fibril and the tropocollagen 5^th^ order collagen peak orientation (Fig. 5A) were homogeneous throughout the leaflet, as expected (Fig. 5B, angle dependency of the orientation encoded in the colour wheel). On the other hand, while the non-calcified explanted leaflet retains to some extent its homogeneous orientation, some areas present low-intensity collagen fibril diameter reflections and a lower degree of orientation for the 5^th^ collagen peak. These changes are visible in the orientation of the bundle structure of the fibirls, the 5^th^ collagen peak (Fig. 5B, parts of the commissures and free edge appear as black), and the scattering pattern (Fig. S12). However, no changes were seen on the collagen nanostructure, as observed from the object orientation in the 0.417 < q < 0.450 nm^-1^ collange/mineral q-equivalent domains (CNQED) (Fig. 5B), suggesting the fibrillary structure with the nanodomain of the tropocollagen molecules remaining unaffected.

Similarly, heterogeneity was observed in the calcified leaflet even in non-mineralised areas (Fig. 5B), with comparable orientations observed for the collagen diameter and the CNQED (shown in the same colour range in Fig. 5B) for mineralised and non-mineralised areas. This suggests that the structures within the calcified regions follow the natural collagen fibril orientation (Fig. 5B and S13) and supports an intra- or inter-fibrillar mineralisation of the collagen structure. Additionally, an altered main orientation of the fibrillar structure within the non to low-calcified regions is associated with a mineralisation fingerprint (visible as a yellow halo around the mineralised part) (Fig. 5B). In these regions, a change in the collagen fibrillar diameter from the healthy diameter calculated to be 144.29 ± 3.09 nm towards 111.16 ± 6.06 nm in the region within the mineralisation front is observed (Fig. S10). Moreover, the characteristic scattering pattern (Fig. S13 and S14) is found to gradually disappear closer to the mineral, suggesting either damages in the tropocollagen helical structure or the formation of an additional overlapping phase suppressing it. Finally, collagen structure changes can be seen from the change in orientation of the 5^th^ collagen peak in regions where the CT does not show any mineralisation (yellow regions on 5^th^ collagen peak orientation) (Fig. 5B). These results illustrate the extent of non-calcific collagen alterations, highlight the impact of implantation on bioprosthetic valvular tissue, and reveal at least a partial connection between such non-calcific alterations and calcific degeneration.

### Macroscopic distribution of haemodynamic and biomechanical indicators correlate with damage

Based on the associations of haemodynamic factors to atheroma plaques in native cardiovascular tissues (54), we hypothesised that similar descriptors may also be employed to predict zones prone to the aforementioned tissue alterations in bioprosthetic valves. We, therefore, explored correlations between macroscopic calcific alterations and specific temporally and spatially varying wall-shear-stress fields using high-fidelity simulations of the coupled blood flow and leaflet mechanics. We investigated the local distribution of biomechanical metrics such as time-averaged wall shear stress (TAWSS), oscillatory shear index (OSI), topological shear variation index (TSVI), scalar strain (SS) and von Mises stress (VMS) using two valvular geometries (ULth0 and Vlth30, Fig. S15). For the ULth0 geometry, we observed high TAWSS in the central parts of the leaflet belly for both sides (Fig. 6A and S16), in line with the tendency of mineral accumulation. However, we also observed higher TAWSS values at the commissures and annulus in the aortic and ventricular sides (Fig. 6A and S16), areas rarely found calcified. These elevated values could result from restricted motion and the narrowness of the flow regions between the leaflets and the stent (Fig. S15) to which the leaflets are attached, promoting more intense shear forces due to blood flow. In line with the regions highlighted by high TAWSS values, the equivalent tensile strength (VMS) in the leaflets is maximum at the level of the commissures and in the middle of the free edge, regions where the scalar strain (SS) is minimum.

**Fig. 6.**
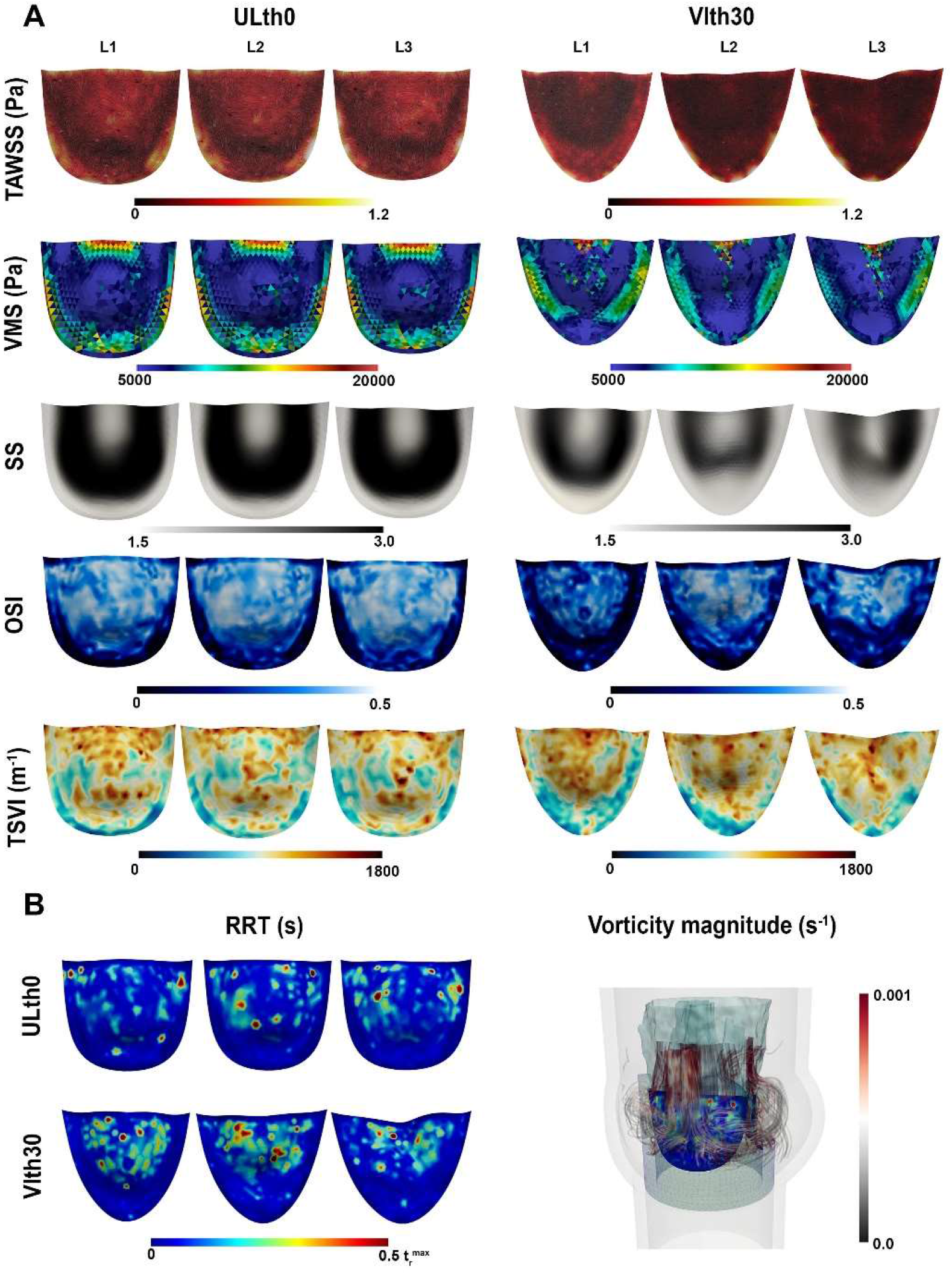
Spatial distribution at the aortic surface of the three leaflets using a fibre-reinforced material model and the leaflet geometries ULth0 and Vlth30. **(A)** Distributions of the time-averaged wall shear stress (TAWSS), von Mises stress (VMS), Scalar strain (SS), oscillatory shear index (OSI) and topological shear variation index (TSVI). **(B)** Relative residence time (RRT) and 3D representation of the streamlines coloured with the vorticity magnitude. The leaflet is colour-coded with the residence time and the isosurface of turbulent kinetic energy (equal to 0.08 m^2^/s^2^) using a fibre-reinforced constitutive law for the leaflet material and a ULth0 and a Vlth30 leaflet geometry. Distributions of the aortic side of the RRT are shown, and the vorticity magnitude of blood in the aortic root (not drawn to scale).

Conversely, high OSI and TSVI values are mainly identified in the centre and top parts of the leaflet belly for both sides (Fig. 6A and S16). These observations are in line with the cumulative increased tendency of mineral accumulation in the centre of the belly in analysed leaflets (Fig. 1C). Subsequently, higher OSI and TSVI could contribute to the triggering of collagen network alterations, leading to subsequent collagen mineralisation; supporting the hypothesis that time-varying haemodynamic wall shear stresses leading to the repetitive expansion and contraction of the leaflet surface may contribute to valvular failure.

The association of blood flow patterns with the presence of minerals on the leaflet surface was also evaluated through computations of the RRT (i.e. the time blood elements spend at a specific location on the surface) and blood vorticity. The RRT distribution (comparably to the TSVI distribution) presents similar patterns on the aortic and ventricular sides, with higher values observed at the top half of the leaflet (Fig. 6B and S16), in line with the similar mineral content (calcified particles and nodules) observed on the aortic and ventricular leaflet external surfaces as identified through the SEM analysis.

The lower vorticity values in the belly region of the leaflets can be explained by the low flow velocities simulated in the central regions of the belly, which are associated with higher residence time (Fig. 6B and S16). The fluid-structure interaction simulations also indicate higher flow vorticity magnitudes near the free edge of the leaflets, which could explain the absence of minerals in this region until later degeneration stages (once the leaflet becomes highly calcified, as observed through microCT imaging). This could be due to the increased flow momentum (related to turbulence) preventing higher mineral deposition, triggered by the generation of vortical structures subsequent to the shear-promoting flutter leaflet motion.

Considering the different leaflet geometries observed in the analysed valves, a different leaflet geometry (Vlth30) (Fig. S15) presenting a distinct bi-directional flutter motion (as compared to that of the ULth0 leaflet geometry) during the peak systolic phase was used to verify that geometry does not alter the overall patterns observed in these indicators (Fig. 6 and S16). Moreover, the Vlth30 leaflet geometry (Fig. S15 and S16) also shed light on a possible link between flutter instability (34, 35) of the leaflets at peak systole and calcification. This possible link is supported by the fact that the high OSI and TSVI regions (in both geometries) seem to be located at the level of the curvature change shown by the high values for scalar strain (Fig. 6 and S16) ensuing from the fluttering leaflet dynamics over peak systole, the region also found to be calcified in all macroscopically calcified leaflets (Fig.6B). The experimental validation using in vitro tomographic particle imaging velocimetry data of the simulation-based flow field quantities (time-averaged and fluctuating velocities) as well as leaflet kinematics (effective orifice area and leaflet shape during opening phase) can be found in Corso et al.(35, 55)

For a more detailed correlation of the location of the macroscopic calcific structures identified through the microCT analysis to the location predicted from the coupled blood-leaflet dynamics simulations, the contributions of different indicators were combined in order to obtain a reconstructed calcification intensity (out of the formulation of an optimisation problem, cf. Eq. 8). This was then directly compared to the observed calcification intensity (calculated from the cumulative spatial distribution of calcification from microCT) in the macroscopically calcified leaflets considered (Fig. 7). The analysis indicates a good agreement between the two calcification patterns (as brought out by the relatively high value of 0.77 for the coefficient of determination R^2^ in the regression) which further supports the possible contribution of these biomechanical indicators and the adequacy of the resulting reconstructed calcification equation for the assessment of the degeneration and calcification of bioprosthetic valves. It is worth noting that the relative contribution of each of the terms in Eq. 8 (when averaged over the 100 bins of the discrete distribution) amounts to 18% for the OSI term, 62% for the TSVI term, the contribution of the shear strain term being negligible. Furthermore, it is interesting to mention that the calcified area estimated from the microCT measurements represents 35% of the total leaflet surface while for the ULth0 leaflet geometry, the prediction from the reconstructed calcification intensity gives an area reaching 23% of the total leaflet surface.

**Fig. 7.**
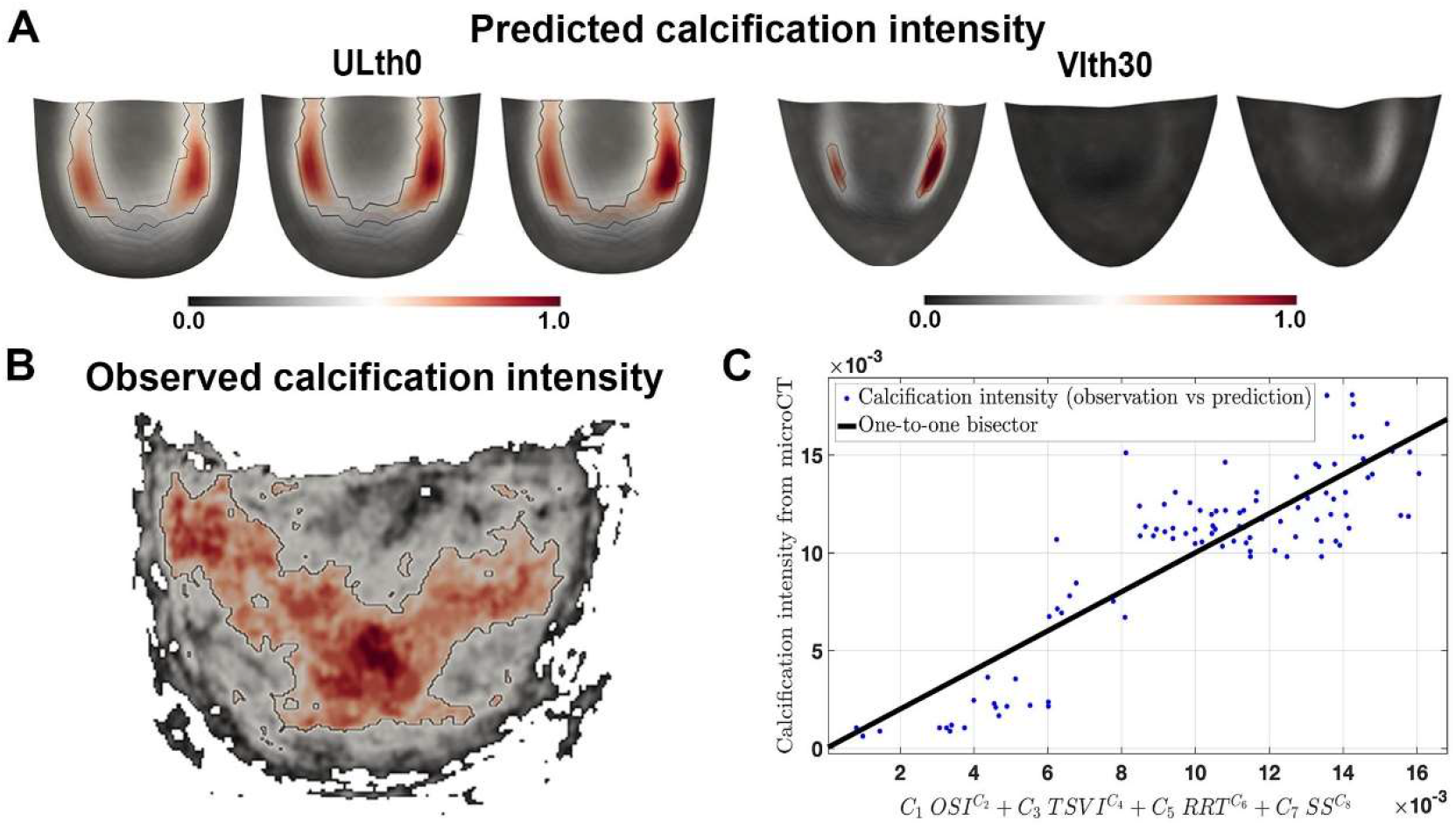
Comparison between predicted calcification intensity and the observed calcification intensity. **(A)** Predicted calcification intensity computed using the contribution of the OSI, TSVI, RRT and SS indicators. **(B)** Observed calcification intensity as observed through the microCT analysis. **(C)** Regression analysis showing a good correlation between the predicted and the observed calcification analysis (R^2^ = 0.77).

## Conclusions

Our multiscale multimodal analysis of bioprosthetic heart valve explants enabled a in-depth, systematic characterisation of minerals with full histoanatomical context and the documentation of the extensive presence of calcified collagen fibres and fibrils within all macroscopically calcified explanted bovine valve leaflets analysed. Even though older transmission electron microscopy studies show collagen mineralisation in failed porcine bioprosthetic valves, and the involvement of collagen in mineralisation has also been discussed in *in vitro* studies on fixed pericardium (56–59), the extent and severity of such calcification in current widely used bovine bioprosthetic valves have not been apparent. Notably, such collagen mineralisation has not been reported in native aortic valves (Fig. S4) or other native soft tissue, or at least not to a comparable extent. This finding suggests an underlying mechanism particularly prevalent in bioprosthetic valves, likely due to controversial tissue pretreatments and fixation, which may be preventable using emerging tissue treatment methods optimized based on nanoanalytical investigations enabling detection of early stage mineralizations. Notably, identifying early-stage mineralisation, such as the ones identified in this work by nanoanalytics, may offer an accelerated route for prescreening alternative tissue treatment options.

Finally, we highlight the importance of considering non-calcific tissue alterations to understand bioprosthetic valve structural failure and their role in mineralisation induction. The discovered disruptions to the collagen network and their link to biomechanical stress have been suggested to be one of the primary reasons limiting durability and playing a role in calcification induction (acting as nucleation sides) (21, 57, 60–62). Interestingly, based on our simulations, we note that the valve with the Vlth30 leaflet geometry is most likely less prone to calcification than the one with the ULth0 leaflet geometry. This observation suggests the existence of calcification-prone valve designs.

Main limitations of the current study include the limited number of explanted valves samples available and the limited patient data. However, the high abundancy of the collagen fiber mineralization (100% of the macroscopically calcified leaflets contained extensive amounts of mineralized fibres) indicates high prevalence of the observed mineral structures, even based on this study population limited in size. Also, the sex of the animal tissue donors remained undisclosed, even though differences in mineral composition and morphology between male and female animals and humans have been described (63). Another limitation of our study is the assumption that the degeneration mechanisms are progressive processes, as analysing the same valve at different time points to gain quantitative data on non-calcific and calcific processes is not technically feasible. Thus, we can only indirectly hint at the anatomical origins of mineral nucleation and collagen network alterations in explanted valves based on the obtained data. However, the correlations between mineral extent and distribution and implantation duration supports the assumption of a progressive process.

Additionally, a limitation in the fluid-structure interaction computational study concerns the reduced time span (0.3 seconds) considered over peak systole, as compared to the time span of the valvular degeneration mechanisms, which can last for 10 to 15 years. In addition, alterations could be induced throughout the cardiac cycle (i.e. during both the systolic and diastolic phases). The fluid fluid-structure interaction stimulation was calculated based on the motion state of leaflets at the systole phase and the stress distribution on leaflets at other moments was not studied. Despite this, trends in the proposed haemodynamic and biomechanical indicators have been identified, and the spatial distribution of a defined calcification intensity observed from microCT measurements has been fitted and convincingly correlated to the distribution of a predicted calcification intensity. Further *in vitro* experimental work building on our observations may be performed to assess the effect of biomechanical forces in the induction of calcific and non-calcific collagen alterations and aid the understanding of the exact processes leading to collagen mineralisation (and which non-calcific alterations enable initial mineral nucleation).

In addition to the above new insights, our work offers a route to seamlessly integrate nanoanalytical characterisation in the clinicopathological analysis to investigate histoanatomical localisation and characteristics of minerals in a wide variety of tissue samples. This route paves the way for larger clinical studies with significantly larger sample numbers, evaluating diagnostic (and/or prognostic) value of mineral characteristics in soft tissues.

## Acknowledgements

We thank the ETH Microscopy Center (ScopeM) (Dr. Anne Greet Bittermann, Dr. Karsten Kunze, Dr. Tobias Schwarz and Justine Kusch-Wieser) and the Empa Electron Microscopy Center for providing access to their microscopes and Magdalena Unterberger for her help in tissue sectioning and staining. The authors acknowledge the Paul Scherrer Institute, Villigen, Switzerland, for the provision of synchrotron radiation beamtime at the beamline cSAXS of the SLS. P.C. would like to thank B. Becsek and F. B. Coulter for the initial sketches and three-dimensional geometrical description of the leaflets, the crown and the aorta that were modified and adapted to perform the FSI simulations. All schematics were created using Biorender.com.

## Author contributions

E.T., S.B., T.C., D.O. and I.K.H conceived the study. E.T. performed experimental microscopy studies, analysed and interpreted data and created the first draft of the manuscript. P.C. performed the FSI computational simulations, analysed and interpreted the computational data, produced the results and figures for the computational part of the study, performed the comparative analysis (microCT vs simulation data), devised and wrote the computational method part of the study, R.Z. performed micro CT scans and 3D analyses, J.A., C.A. and M.L performed SAXS synchrotron measurements, analysed the SAXS data and wrote the SAXS method part, P.P.H. collected samples and patient data. All authors discussed the data and edited the manuscript. T.C. supervised the clinical part, D.O. the simulation work and I.K.H. the multiscale characterisation work and the overall study.

## Data and materials availability

All data and materials needed to evaluate the conclusions of this paper are present in the main text or supplementary materials. Processable data files can be obtained from the authors.

