## Supplementary Information for "Multiscale Multimodal Characterization and Simulation of Structural Alterations in Failed Bioprosthetic Heart Valves"

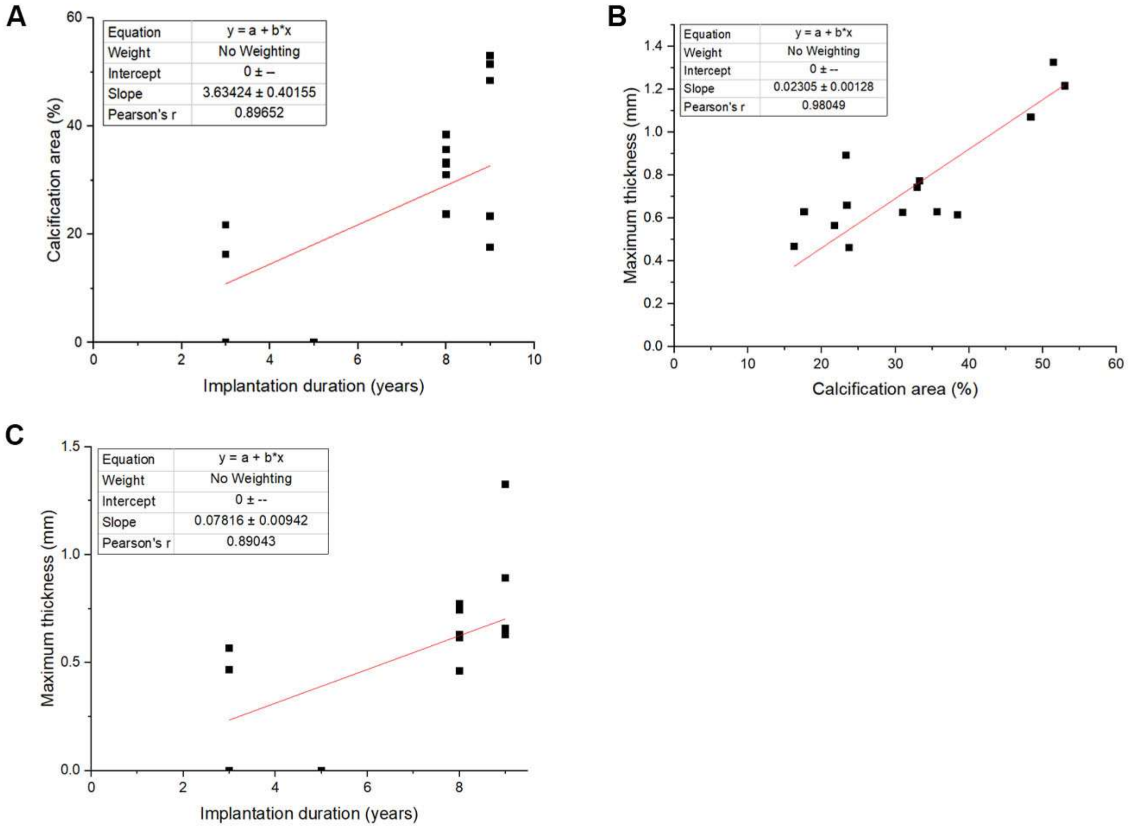


*Fig. S1.* ***(A)*** *Relationship between the calcification area percentage and implantation duration (N=21, Pearson's r: 0.90).* ***(B)*** *Relationship between maximum thickness and calcification area percentage was observed (n = 14, Pearson's r: 0.98).* ***(C)*** *Relationship between maximum thickness and implantation duration (N = 21, Pearson's r: 0.89).*

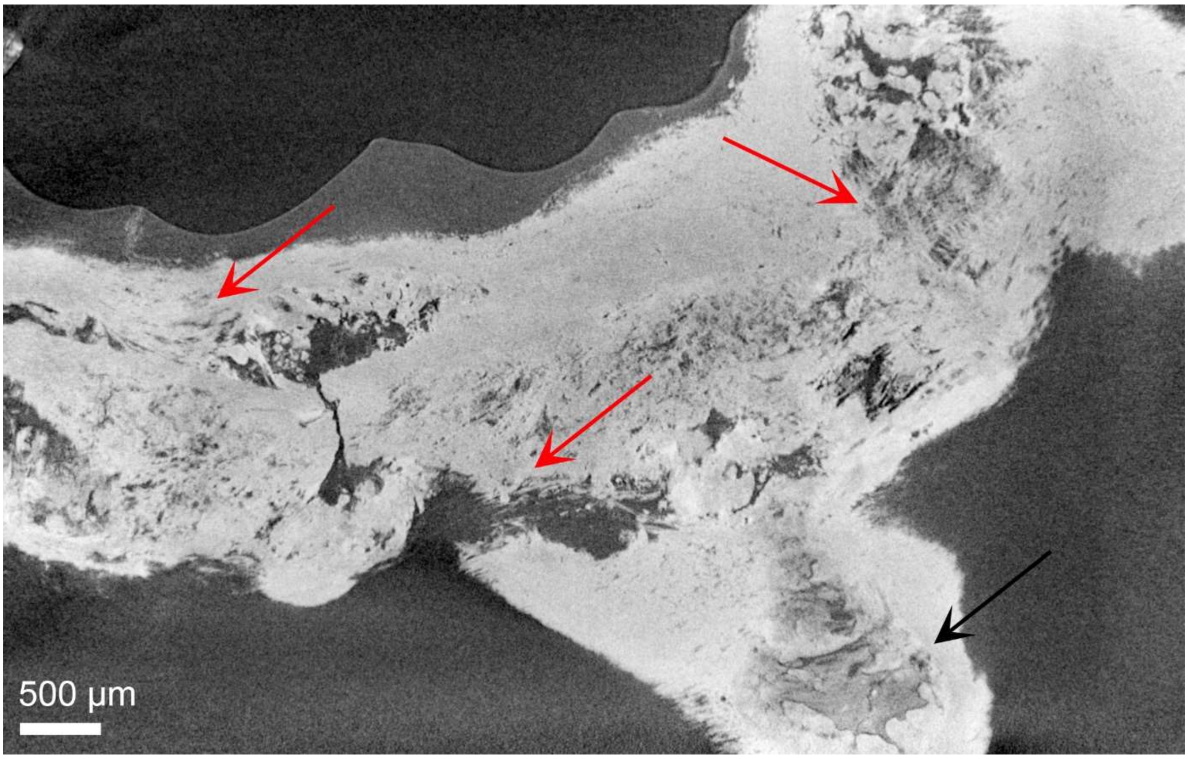


*Fig. S2. High-resolution local microCT cross-section of a moderately calcified bioprosthetic valvular leaflet showing the presence of a compact mineral of no specific morphology (black arrow) and fibrous structures (red arrows).*

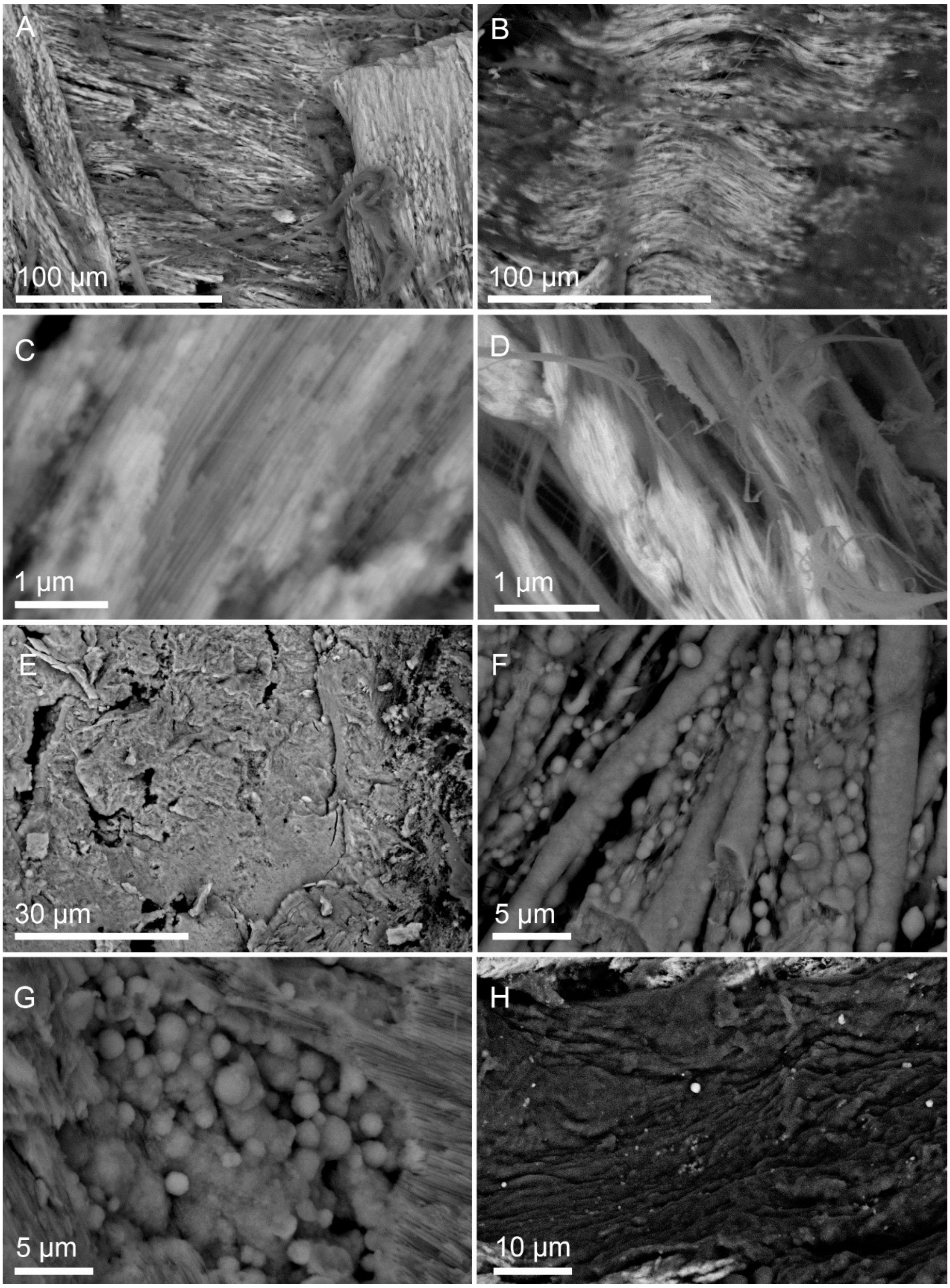


*Fig. S3. Backscattered electron images taken from bioprosthetic leaflets showing **(A, B)** calcified fibres. **(C, D)** calcified fibrils, **(E)** compact calcification **(F, G),** occasionally highly calcified fibres and calcified particles merged, resembling calcified rods and pearl string structures in some cases. **(H)** Calcified particles were also detected as individual entities in tissue parts where macrocalcifications were absent.*

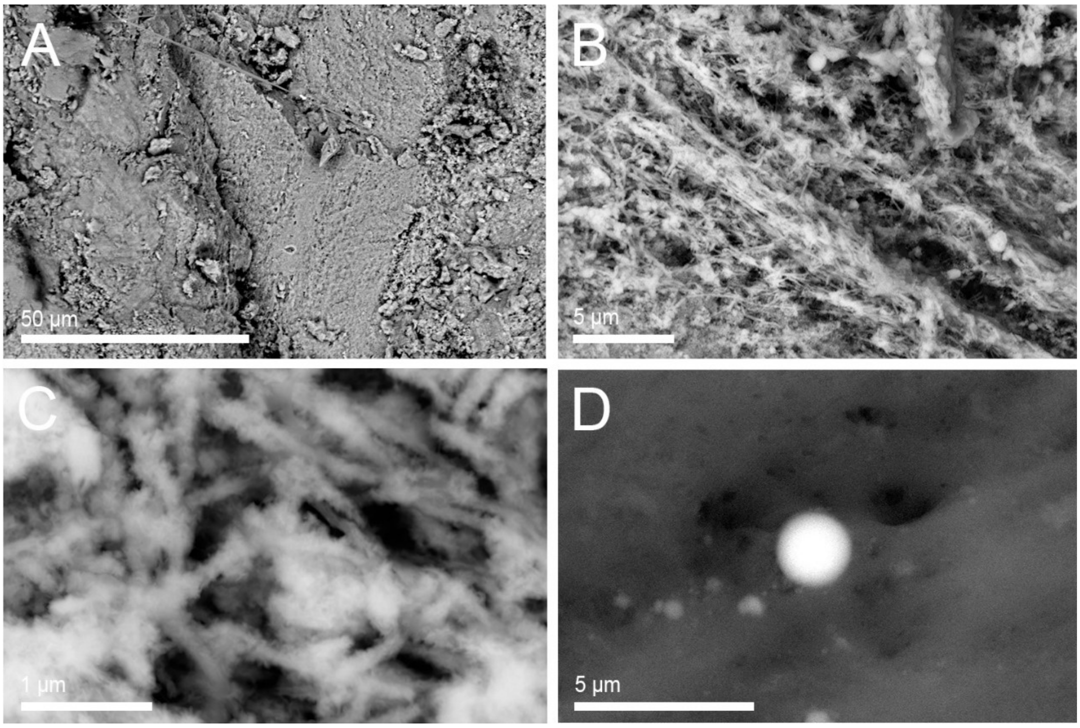


*Fig. S4. Representative backscattered images of minerals found in native valves.* ***(A)*** *Large compact calcifications,* ***(B)*** *fibrous structures made of* ***(C)*** *unorganised needle-like structures and* ***(D)*** *calcified spherical particles*

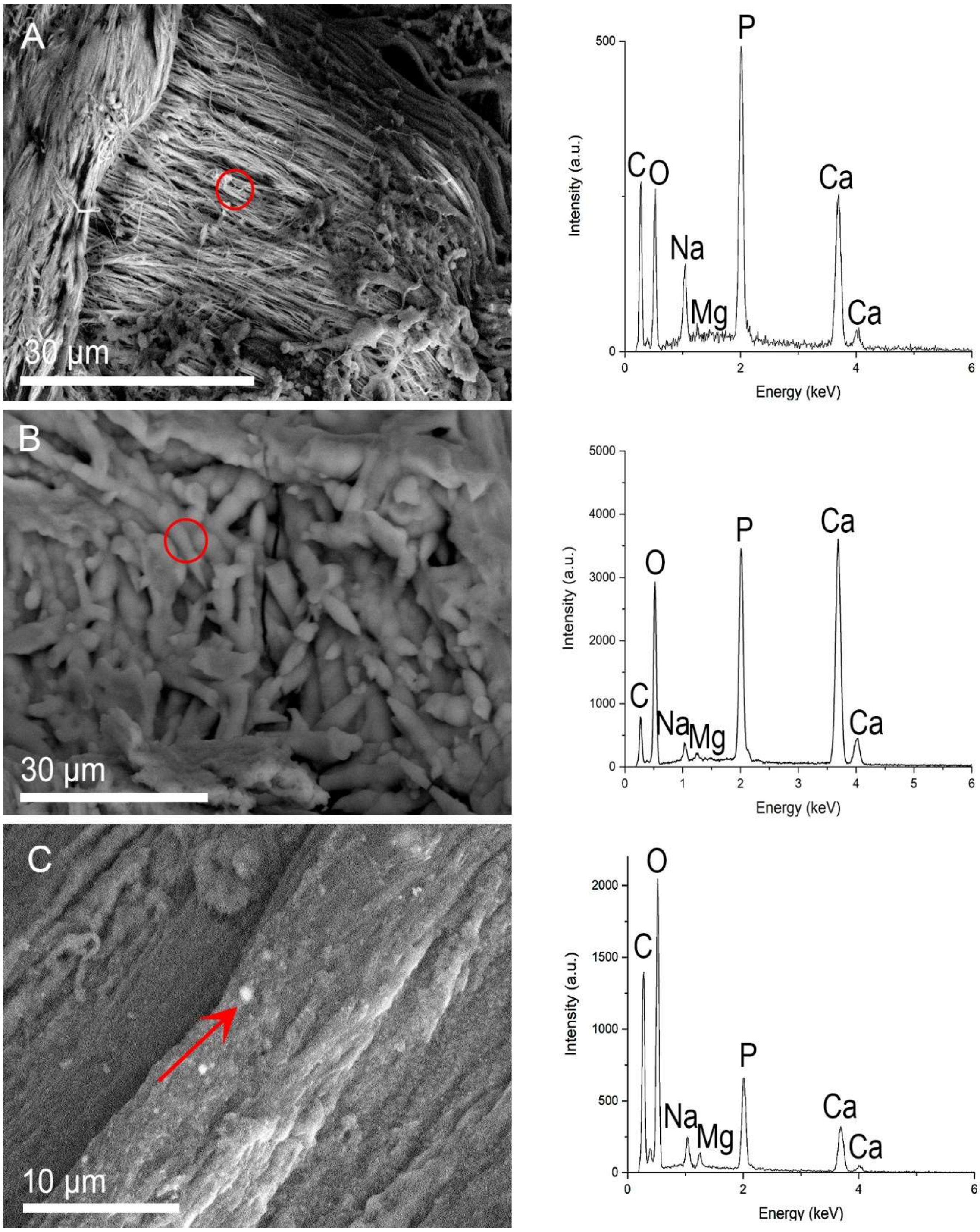


*Fig. S5. Backscattered electron images and corresponding EDX spectra of* ***(A)*** *calcified fibres,* ***(B)*** *calcified needle-like structures and* ***(C)*** *calcified particles observed in macroscopically calcified bioprosthetic leaflets showing a chemical composition of calcium, phosphorus and magnesium. Sodium peaks were also identified, originating from the tissue. The EDX spectra were acquired from the area marked by a circle in* ***A*** *and* ***B****, and by an arrow in* ***C.***

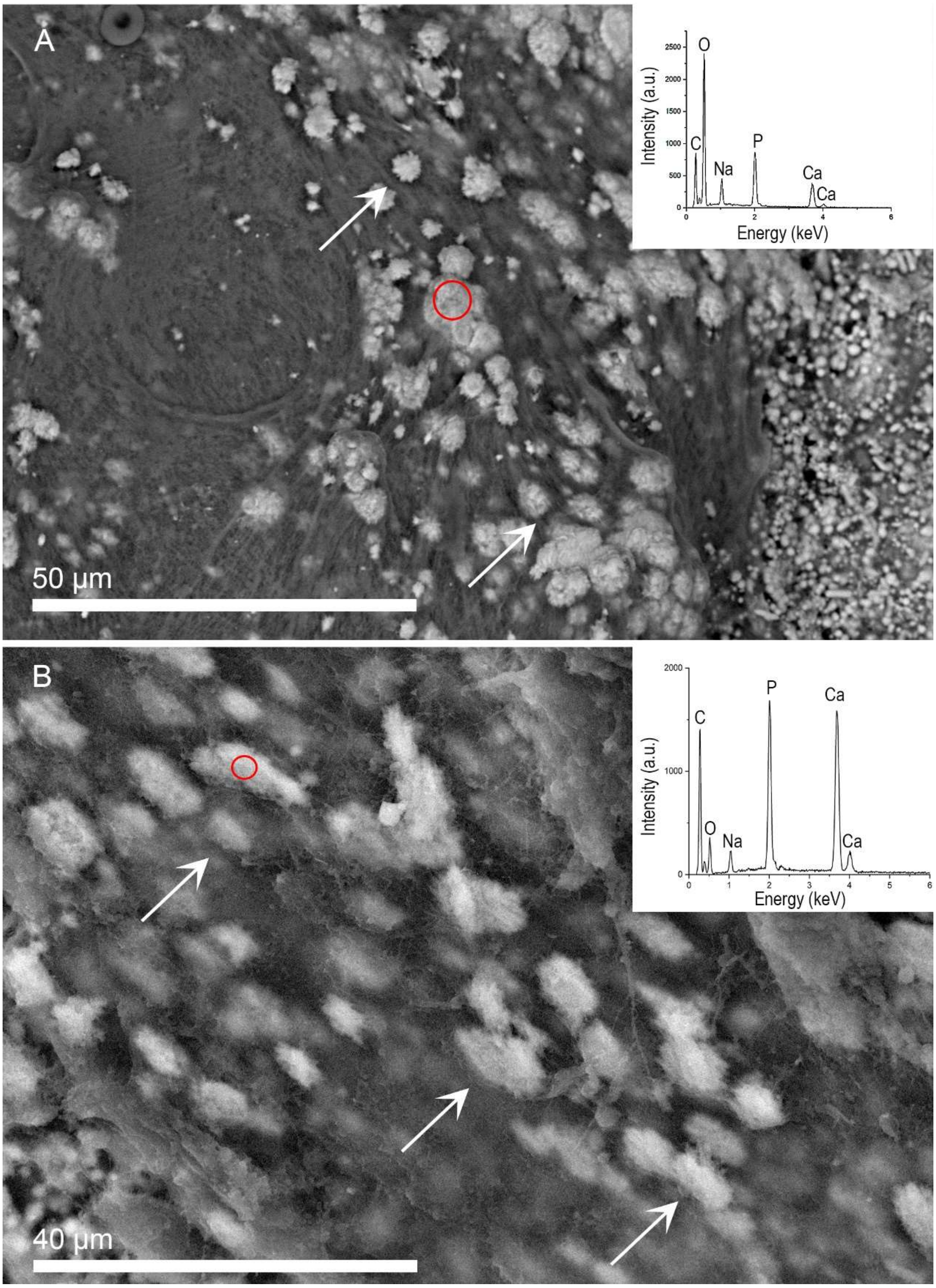


*Fig. S6. Backscattered electron images of calcified nodules (arrows) observed on the* ***(A)*** *ventricular surface of a highly calcified leaflet* ***(B)*** *aortic side of a moderately calcified leaflet along with EDX spectra (insets) of the areas marked with red circles showing a chemical composition of calcium and phosphorus.*

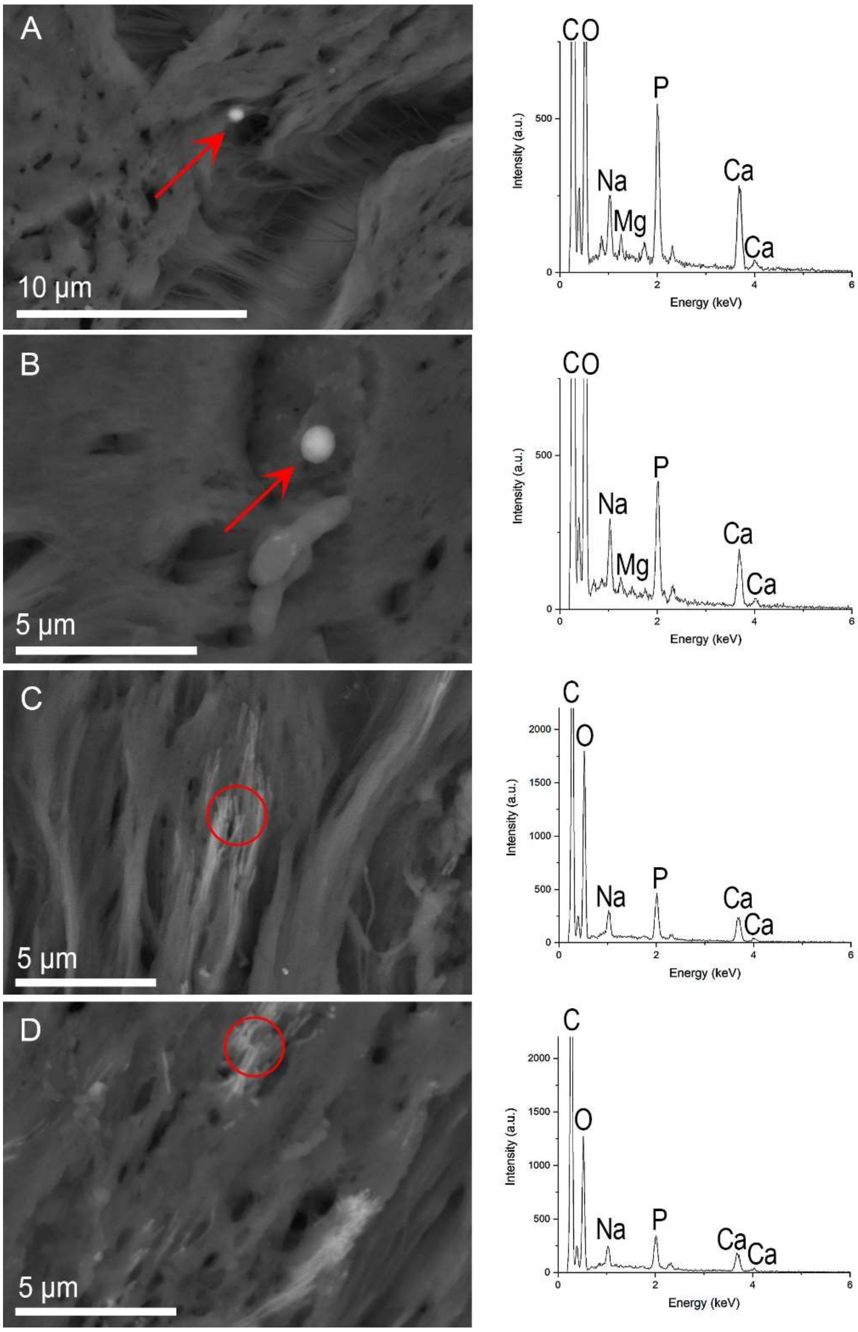


*Fig. S7. Backscattered electron images and EDX spectra of* ***(A, B)*** *calcified particles and* ***(C, D)*** *calcified fibrils of histological slides from non-macroscopically calcified leaflets, showing that the calcified particles are composed of calcium, phosphorus and magnesium, while the calcified fibrils of calcium and phosphorus. EDX spectra were acquired from regions indicated by arrows in* ***A*** *and* ***B****, and circles in* ***C*** *and* ***D.*** *Peaks from sodium and sulphur originated from the tissue and zinc and silicon from the glass background originating from the histological slides the tissue was deposited on.*

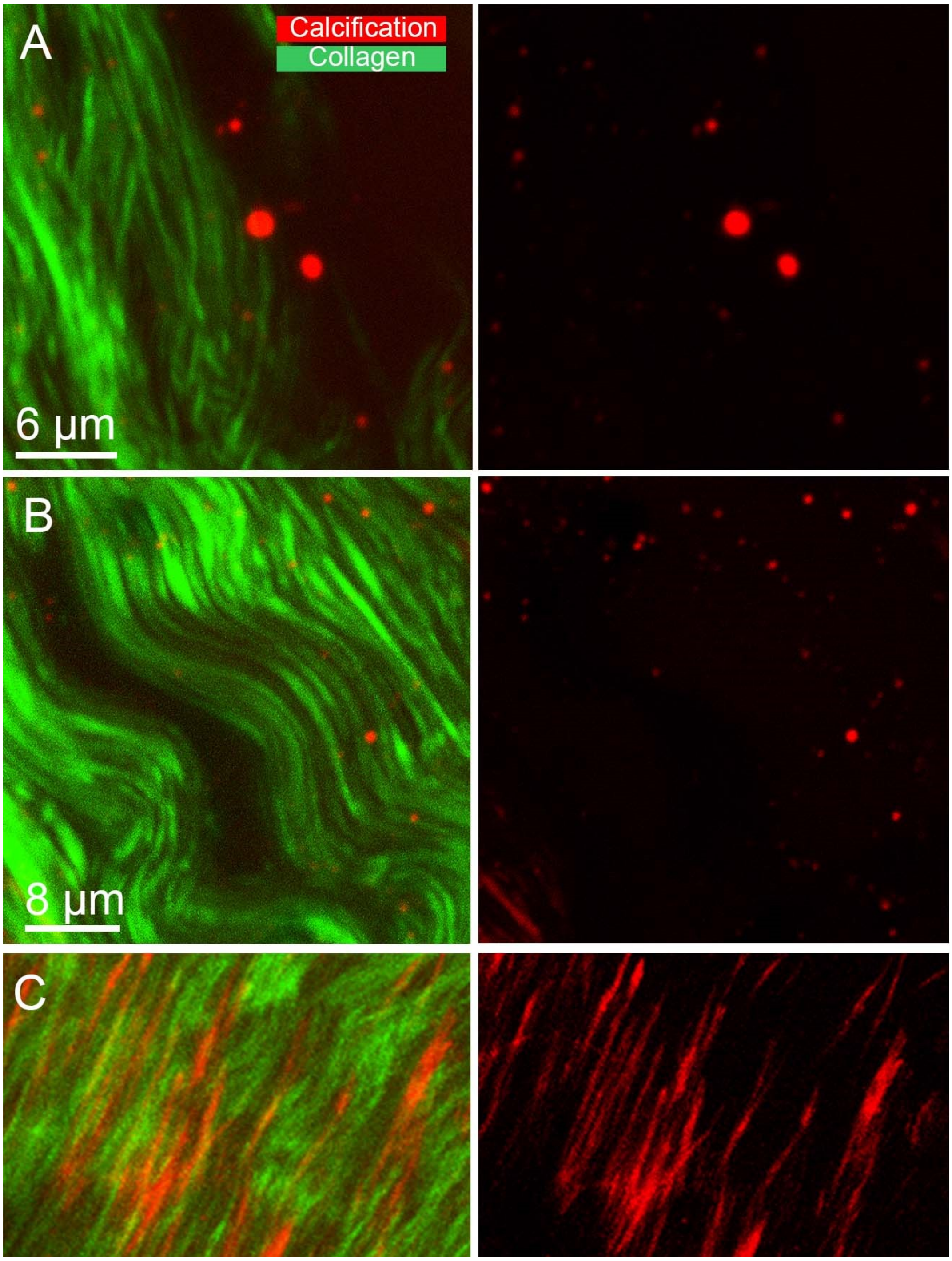


*Fig. S8. Multiphoton micrographs showing **(A, B)** Calcified particles found in gaps between collagen fibres and **(C, D)** calcified collagen fibres.*

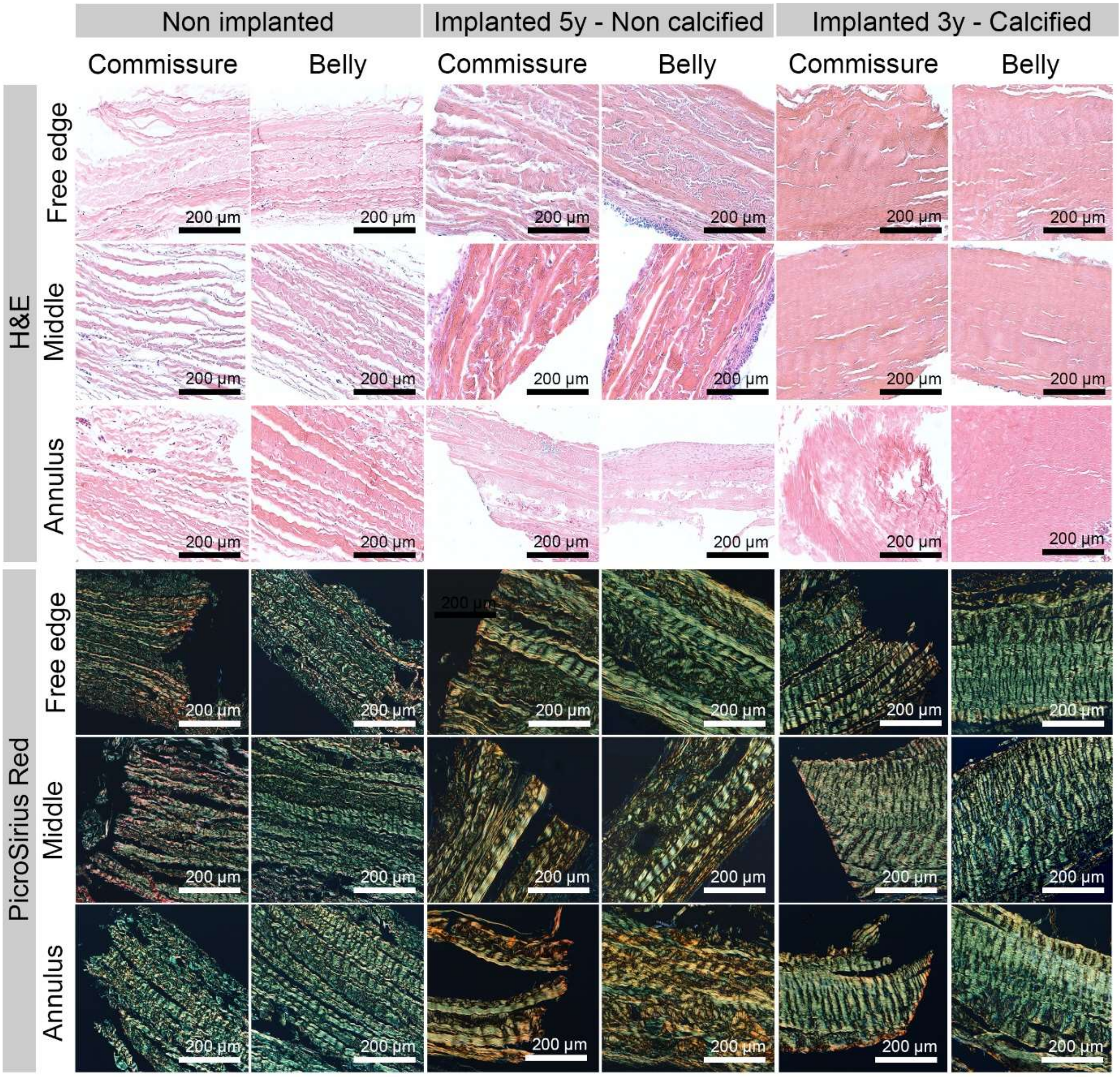


*Fig. S9. Images from H&E and PicroSirius Red stained sections of non-implanted, non-calcified implanted and calcified implanted leaflets acquired from different anatomical locations.*

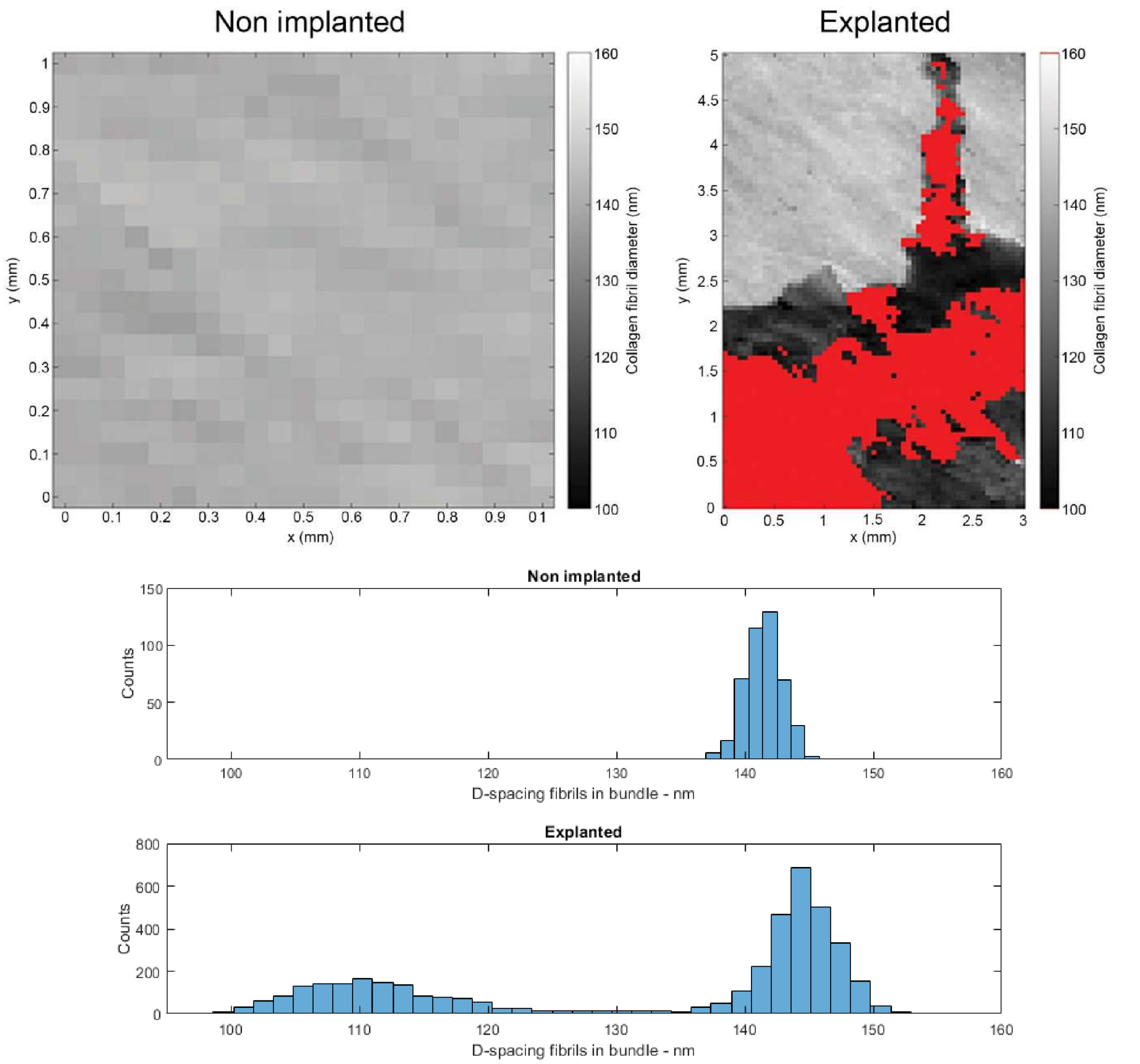


*Fig. S10. Collagen fibril diameter in the non-implanted and explanted samples. The red region indicates a highly mineralised region where no information on d-spacing of fibres in bundles peak was extractable. Histogram of d-spacing sizes over the image. Distribution were fitted with a Gaussian curve, and a peak maxima of 141.46 ± 1.44 nm was found for the non implanted sample. These maxima shifted to 144.29 ± 3.09 nm and 111.16 nm ± 6.06 for healthy and partially calcified regions of the explanted sample, respectively.*

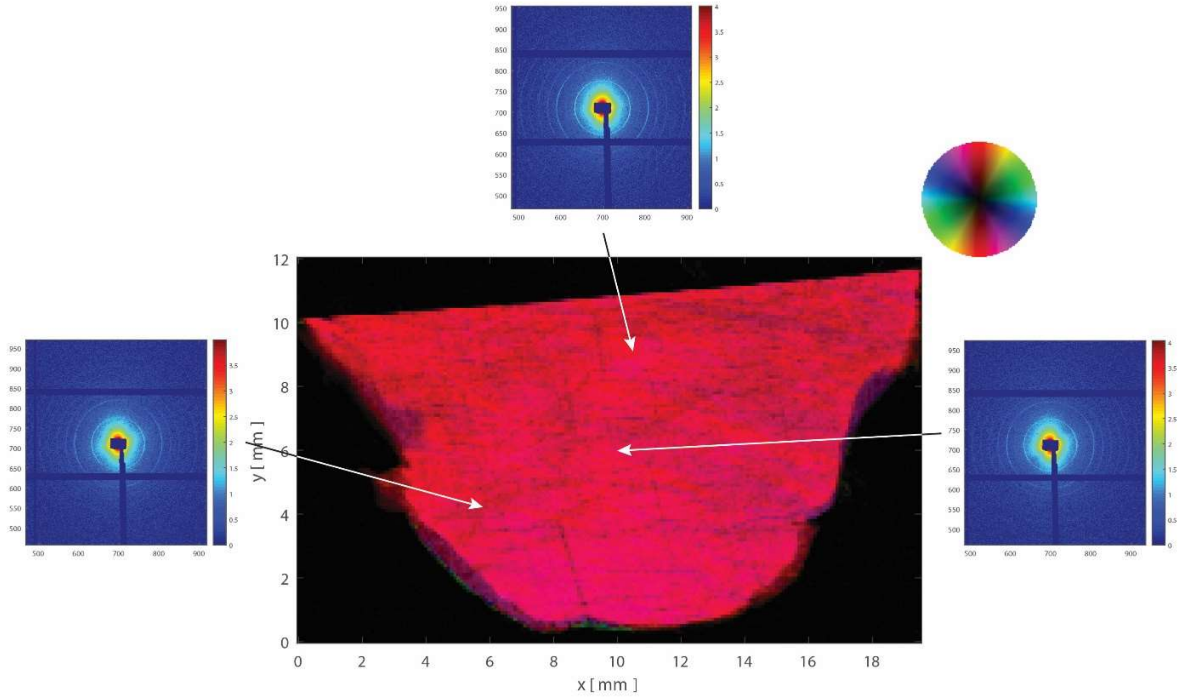


*Fig S11. Representative scattering patterns of collagen observed in a non-implanted leaflet, where the typical 'eye brow' pattern, known to originate from the triple helix structure of the collagen fibrils is uniformly observed throughout the sample.*

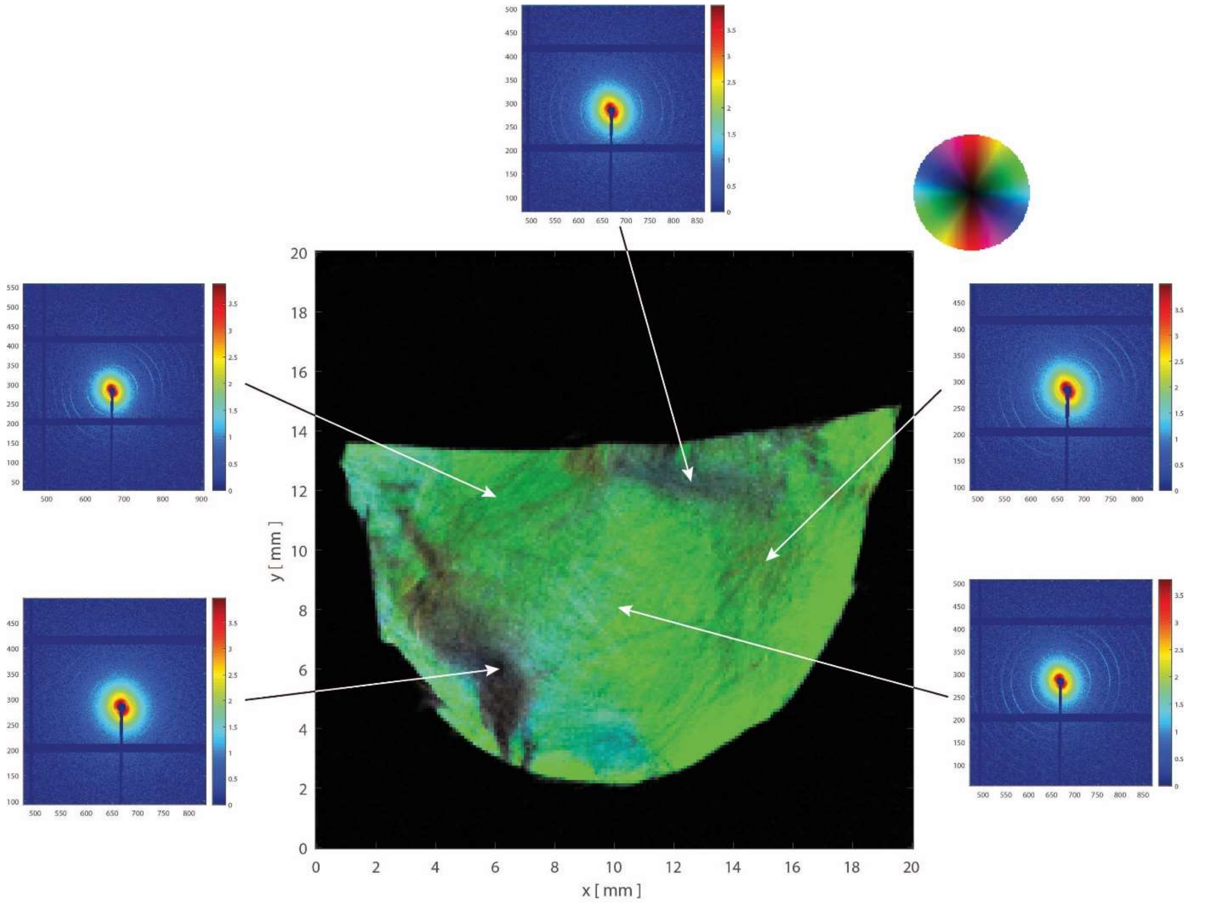


*Fig. S12. Representative scattering patterns of collagen observed in an explanted non-calcified leaflet, where the typical 'eye brow' pattern, known to originate from the collagen fibrils' triple helix structure, is uniformly observed in most of the sample. The scattering pattern was altered in regions where collagen disturbances in the 5th order collagen peak were observed (dark regions).*

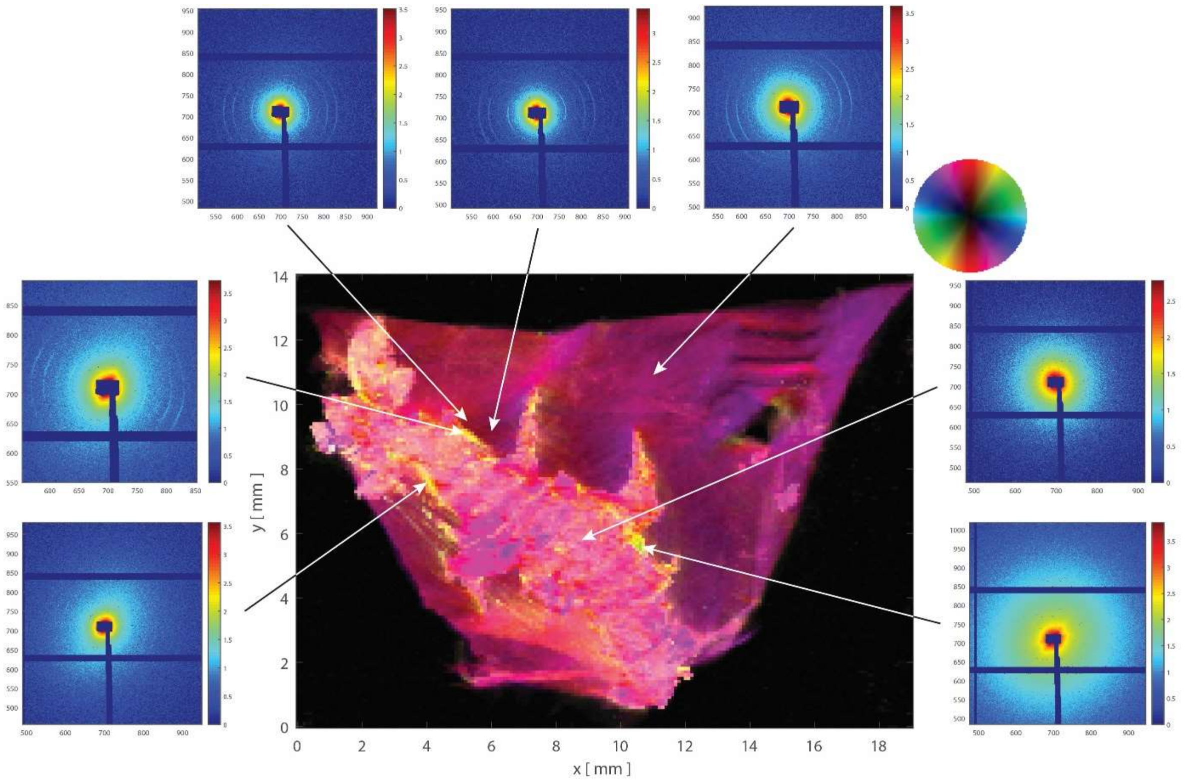


*Fig. S13. Representative scattering patterns of collagen observed in an explanted calcified leaflet, where the typical 'eye brow' pattern can only be observed in non-calcified regions while it is no longer observable in regions at the edges and within the mineral. Additionally, no higher order peak of the collagen can be found in the mineral phase suggesting a denaturation of the collagen structure by the mineral phase.*

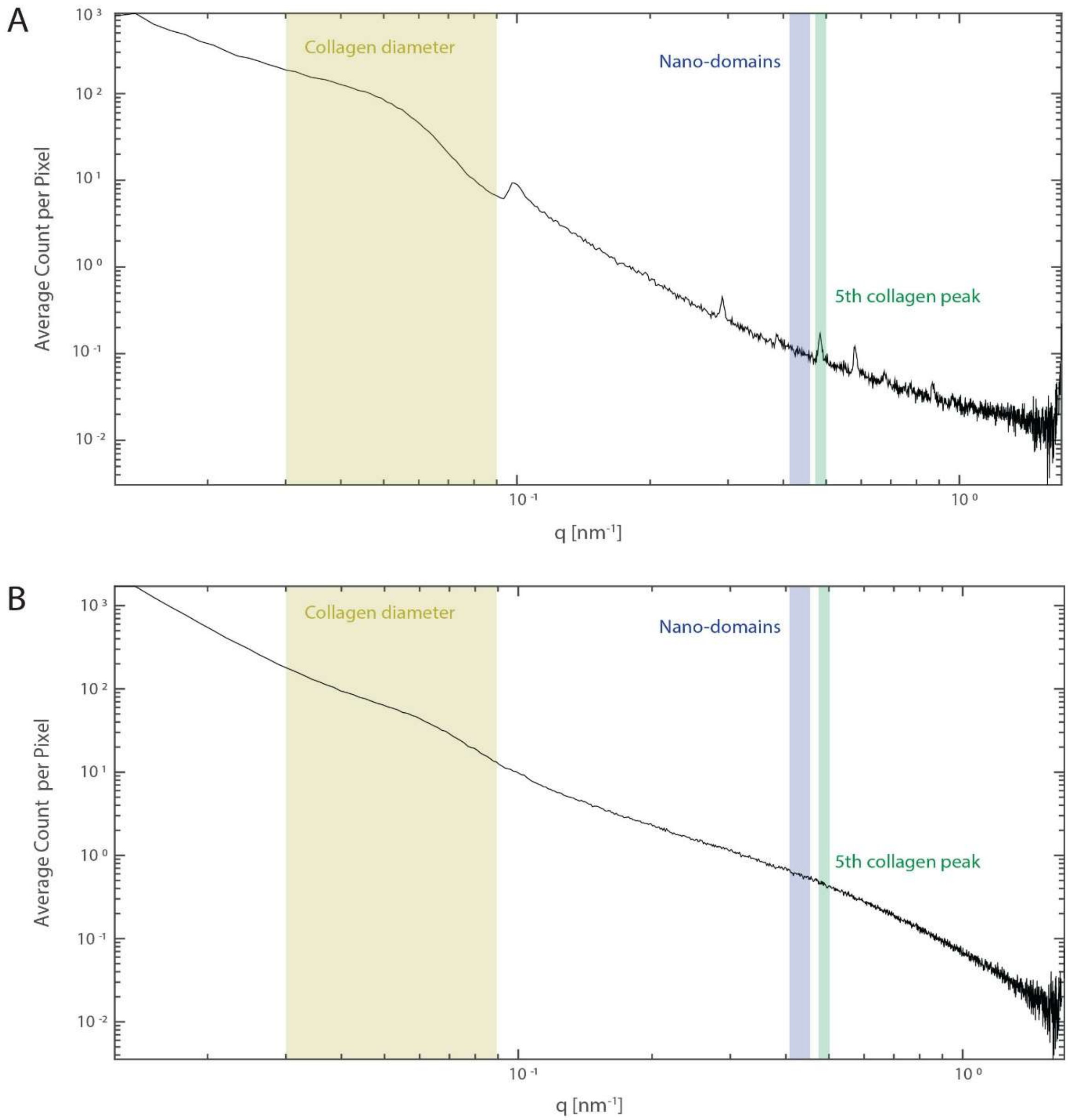


*Fig. S14. Diffraction peak plots showing the q values used for the collagen diameter, nanodomain and 5$^{th}$ collagen peak analysis.* ***(A)*** *Typical graph obtained from collagenous region.* ***(B)*** *Typical graph obtained from a calcified region where most collagen related peaks disappear.*

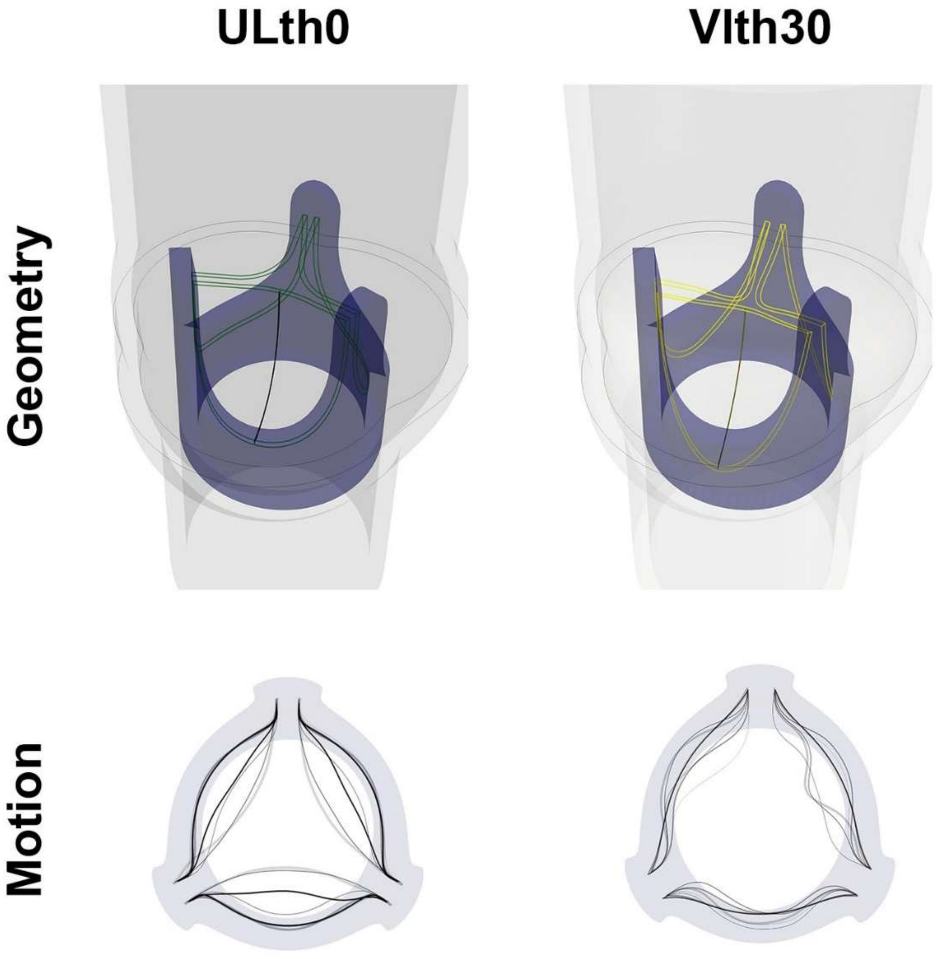


*Fig. S15. Leaflet geometry presented along with the stent and the aorta and motion simulated from the coupled haemodynamics and leaflet dynamics simulations for the geometries ULth0 and Vlth30.*

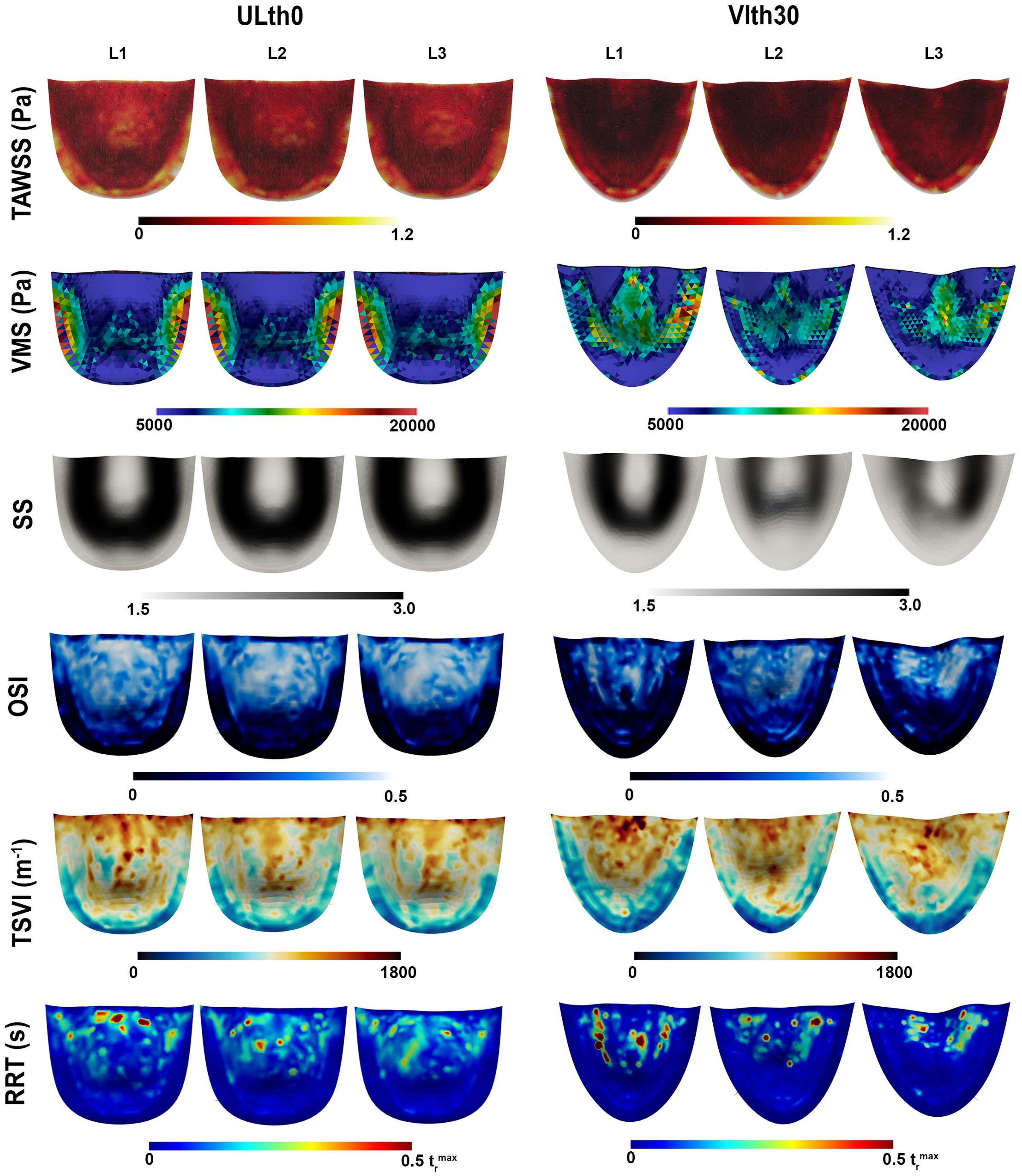


*Fig. S16. Spatial distribution at the ventricular surface of the three leaflets using a fibre-reinforced material model and the leaflet geometries ULth0 and Vlth30. Distribution the time-averaged wall shear stress (TAWSS), von Mises stress (VMS), scalar strain (SS), oscillatory shear index (OSI), topological shear variation index (TSVI) and relative residence time (RRT) are shown.*

*Table S1: Specifications of bioprosthetic valves obtained and individual leaflets analysed.*

| **Valve number** | **Type** | **Implantation duration** | **Leaflets analysed** | |
|---|---|---|---|---|
| | | | **Leaflet** | **Degree of calcification based on microCT<br>(% of calcified area)** |
| 1 | Perimount Magna Ease 23mm | 5 years | 1-1 | Non calcified |
| | | | 1-2 | Non calcified |
| | | | 1-3 | Non calcified |
| 2 | Perimount Magna Ease 23mm | 9 years | 2-1 | Calcified (53%) |
| | | | 2-2 | Calcified (51%) |
| | | | 2-3 | Calcified (48%) |
| 3 | Perimount Magna Ease 23mm | 3 years | 3-1 | Non calcified |
| | | | 3-2 | Calcified (22%) |
| | | | 3-3 | Calcified (16%) |
| 4 | Perimount Magna Ease 25mm | 8 years | 4-1 | Calcified (36%) |
| | | | 4-2 | Calcified (24%) |
| | | | 4-3 | Calcified (33%) |
| 5 | Perimount Magna Ease 23mm | 3 years | 5-1 | Non calcified |
| | | | 5-2 | Non calcified |
| | | | 5-3 | Non calcified |
| 6 | Perimount Magna Ease 25mm | 8 years | 6-1 | Calcified (38%) |
| | | | 6-2 | Calcified (31%) |
| | | | 6-3 | Calcified (33%) |
| 7 | Perimount Magna Ease 25mm | 9 years | 7-1 | Calcified (18%) |
| | | | 7-2 | Calcified (23%) |
| | | | 7-3 | Calcified (23%) |
| 8 | | Non-implanted (new) | 8-1 | Non calcified |

| | | | | |
|---|---|---|---|---|
| | Perimount Magna EASE 23 mm | | 8-2 | Non calcified |
| | | | 8-3 | Non calcified |
| 9 | Perimount Magna EASE 23 mm | Non-implanted (expired) | 9-1 | Non calcified |
| | | | 9-2 | Non calcified |
| | | | 9-3 | Non calcified |
| 10 | Perimount Magna EASE 21 mm | Non-implanted (expired) | 10-1 | Non calcified |
| | | | 10-2 | Non calcified |
| | | | 10-3 | Non calcified |
